# Single-cell Transcriptome Mapping Identifies Common and Cell-type Specific Genes Affected by Acute Delta9-tetrahydrocannabinol in Humans

**DOI:** 10.1101/638254

**Authors:** Ying Hu, Mohini Ranganathan, Chang Shu, Xiaoyu Liang, Suhas Ganesh, Chunhua Yan, Xinyu Zhang, Bradley E Aouizerat, John H Krystal, Deepak C. D’Souza, Ke Xu

**Affiliations:** Center for Biomedical Information and Information Technology, National Cancer Institute, MD, U.S.A.; Department of Psychiatry, Yale School of Medicine, New Haven, CT, 06516; Connecticut Veteran Healthcare System, West Haven, CT, 06515; Bluestone Center for Clinical Research, College of Dentistry, New York University, NY, 10010; Department of Oral and Maxillofacial Surgery, College of Dentistry, New York University, NY, 10010

**Author notes:** Please direct all correspondence to: Ke Xu, MD, PhD, Associate Professor of Psychiatry, Yale School of Medicine.

**Keywords:** Delta-9-tetrahydrocannabinol, single cell transcriptome, peripheral blood mononuclear cells, differential gene expression, gene set enrichment analysis, gene co-expression

## Abstract

Delta 9-tetrahydrocannabinol (THC), the principal psychoactive constituent of cannabis, is also known to modulate immune response in peripheral cells. The mechanisms of THC’s effects on gene expression in human immune cells remains poorly understood. Combining a within-subject design with single cell transcriptome mapping, we report that administration of THC acutely alters gene expression in 15,973 human blood immune cells. Controlled for high inter-individual transcriptomic variability, we identified 294 transcriptome-wide significant genes among eight cell types including 69 common genes and 225 cell-type specific genes affected by acute THC administration, including those genes involving not only in immune response, cytokine production, but signal transduction, and cell proliferation and apoptosis. We revealed distinct transcriptomic sub-clusters affected by THC in major immune cell types where THC perturbed cell type-specific intracellular gene expression correlations. Gene set enrichment analysis further supports the findings of THC’s common and cell-type specific effects on immune response and cell toxicity. We found that THC alters the correlation of cannabinoid receptor gene, *CNR2*, with other genes in B cells, in which *CNR2* showed the highest level of expression. This comprehensive cell-specific transcriptomic profiling identified novel genes regulated by THC and provides important insights into THC’s acute effects on immune function that may have important medical implications.

## Main

With the increasing rates of cannabis use for recreational and medical purposes, it is important to address a gap in our understanding of its impact on immune and inflammatory functions.^1, 2^ Delta-9-tetrahydrocannabinol (THC), the principal psychoactive constituent of cannabis, powerfully modulates immune function in peripheral cells,^3^ in part, through activating cannabinoid receptor 2 (CBR2).^1, 3–6^ *In vitro* studies of cannabis exposure, which contains over 450 compounds, show that it modulates immune function,^7–9^ changes cytokine production,^8, 10, 11^ inhibits cell proliferation,^2^ and induces apoptosis.^12, 13^ However, little is known about the mechanisms of *in vivo* THC exposure on the transcriptomes of distinct types of peripheral blood mononuclear cells (PBMCs) in humans.

Single cell RNA-seq (scRNA-seq) offers an unprecedented resolution to detect drug effects on cell-specific gene expression in an unbiased fashion^14, 15^ and enables the evaluation of molecular aspects of immune cell heterogeneity.^16^ Few studies have applied scRNA-seq to detect differentially expressed genes (DEGs) induced by drug exposure, and none have evaluated the effects of THC in humans. This limitation is due mostly to high inter-individual transcriptomic variability and types of cells that confound the assessment of the impact of environmental factors. Most recently, a scRNA-seq study identified a large number of common and cell-type specific DEGs for Alzheimer disease, suggesting the improvement of analytical methods to overcome the challenge of high transcriptomic variability.^17^ Here, we report a first sc-RNA seq study using within-subject combined with linear mixed model (LMM) analysis to detect genes affected by THC at single cell resolution.

In this study, samples of blood were drawn and PMBCs extracted prior to (pre-THC) and 70 minutes following (post-THC) a single 0.03mg/kg intravenous dose of THC in two healthy individuals. The selected THC dose reliably produces effects consistent with cannabis intoxication.^18, 19^ The timing of the blood sampling was selected to maximize the likelihood of detecting changes in drug-induced gene expression. A battery of subjective and cognitive assessments were administered to capture the effects and safety of THC.^18, 20, 21^ We profiled the four PBMC samples (two pre-THC and two post-THC) on the 10X Genomics platform.^22^ Quality control processing yielded a total of 15,973 cells and 21,430 genes for analyses (**Figure 1a**).

**Figure 1.**
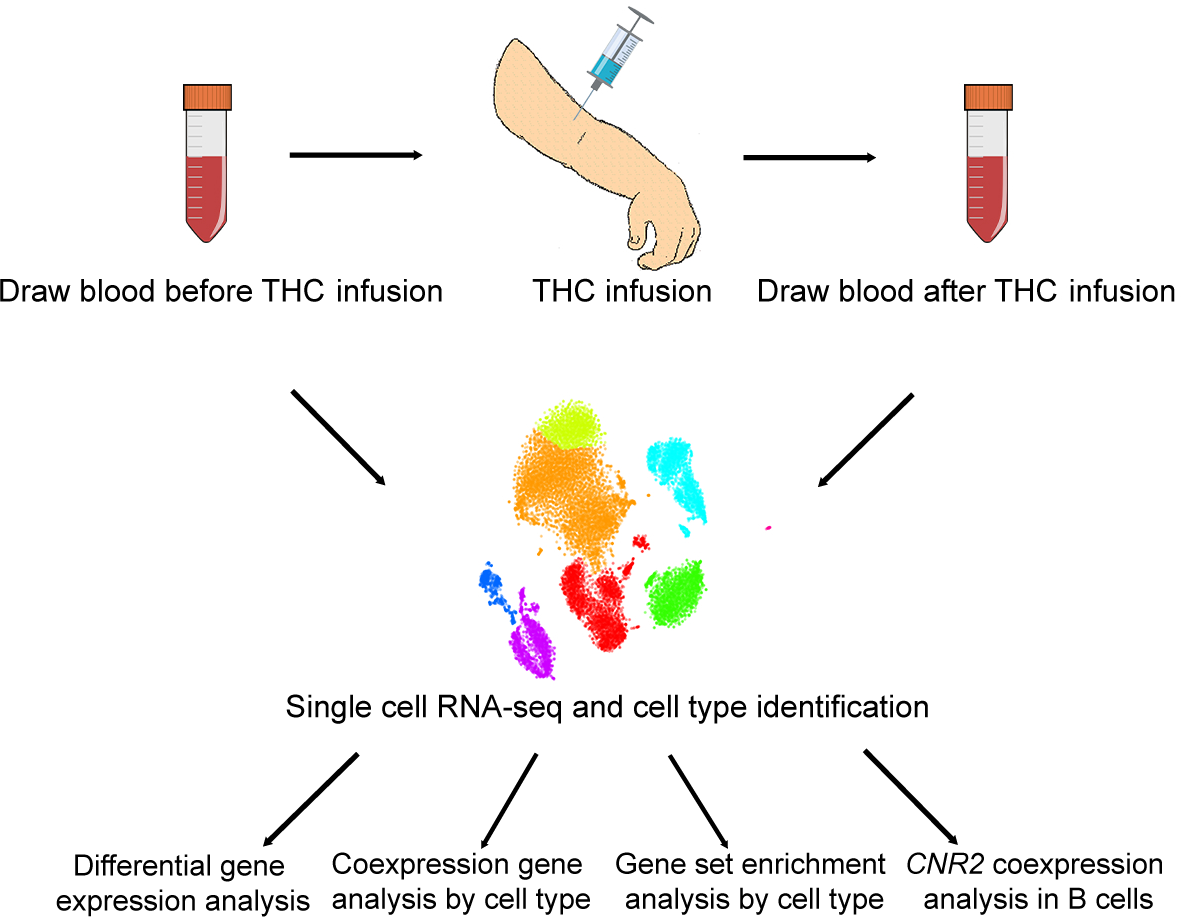
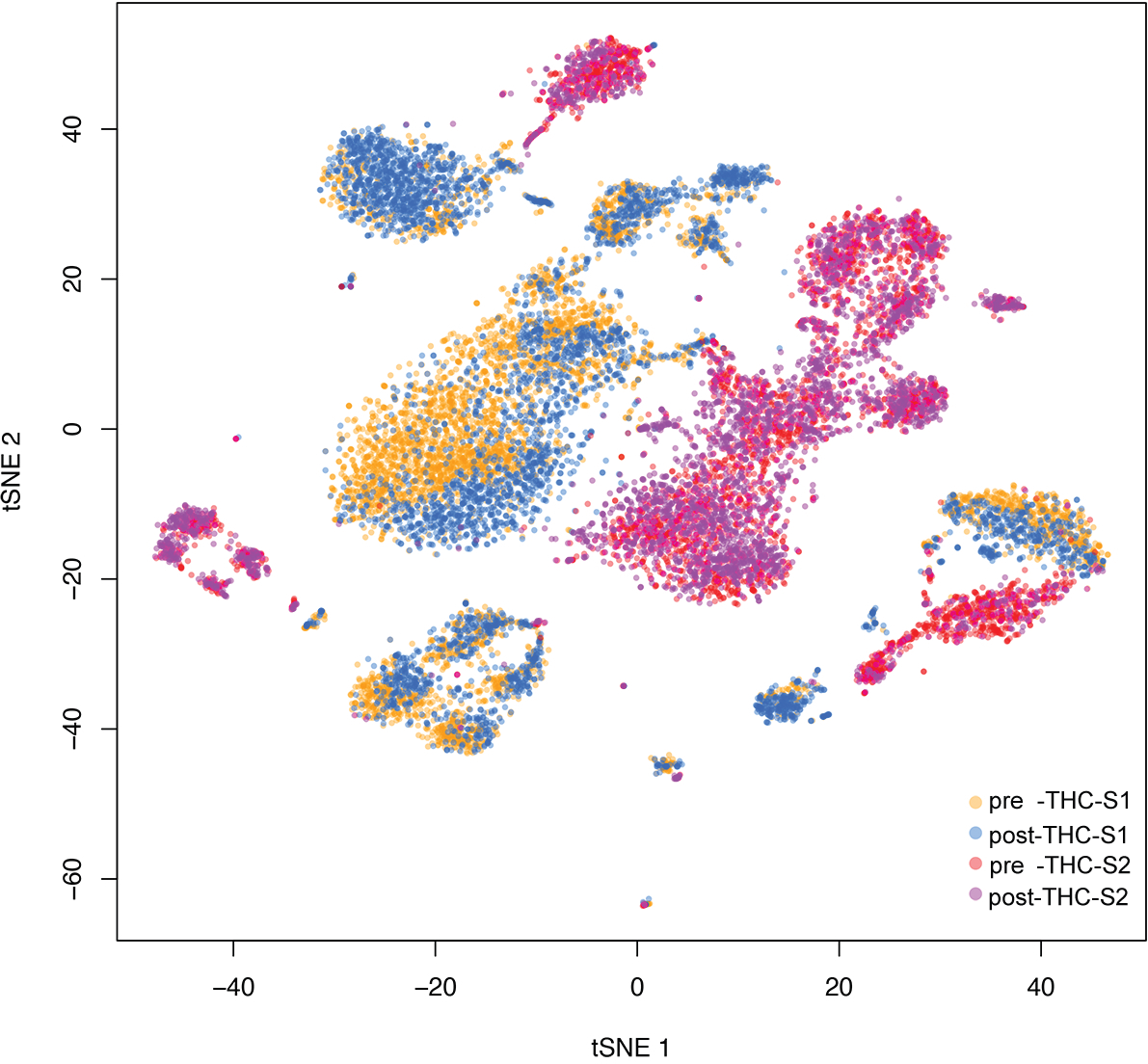
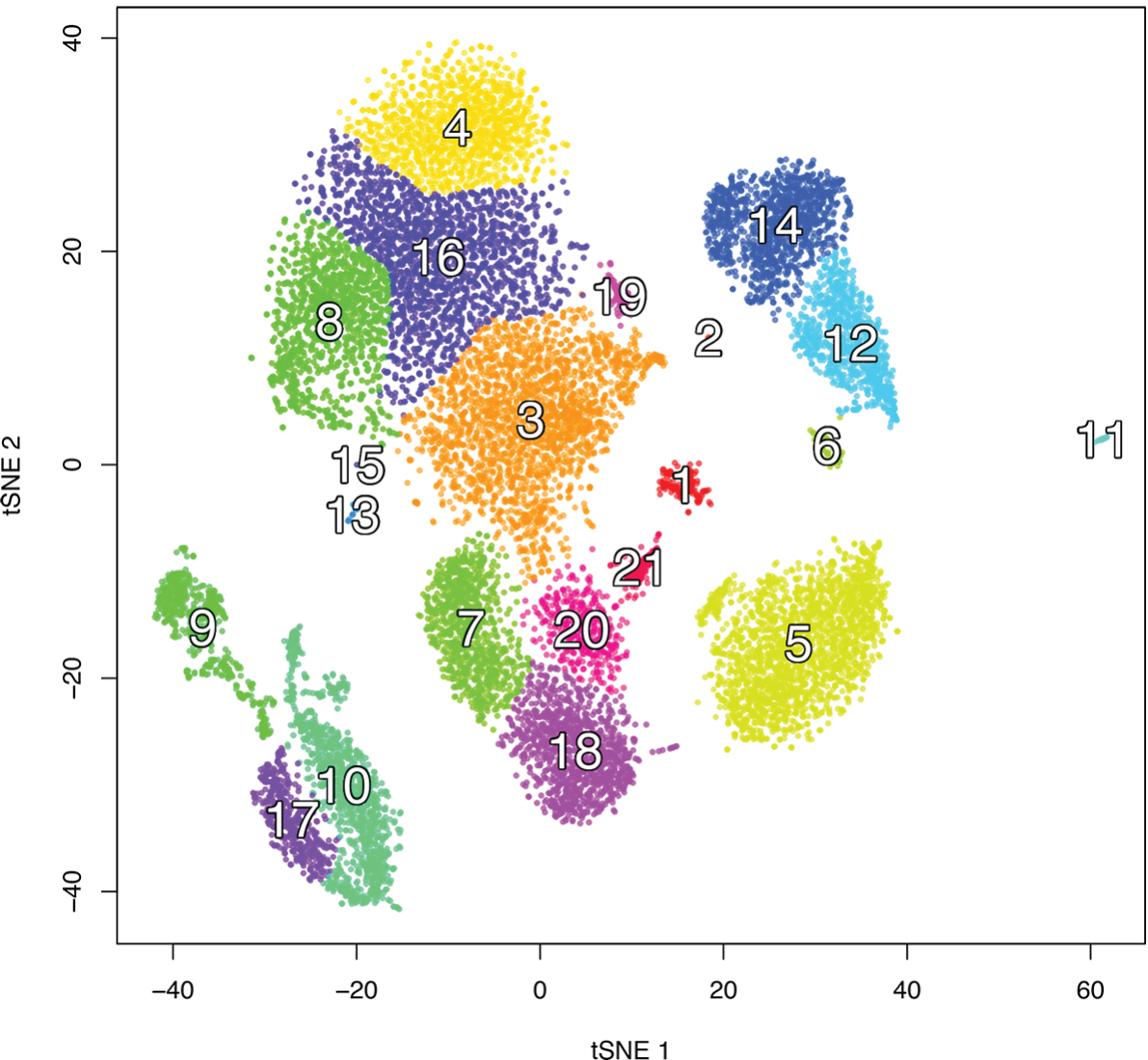
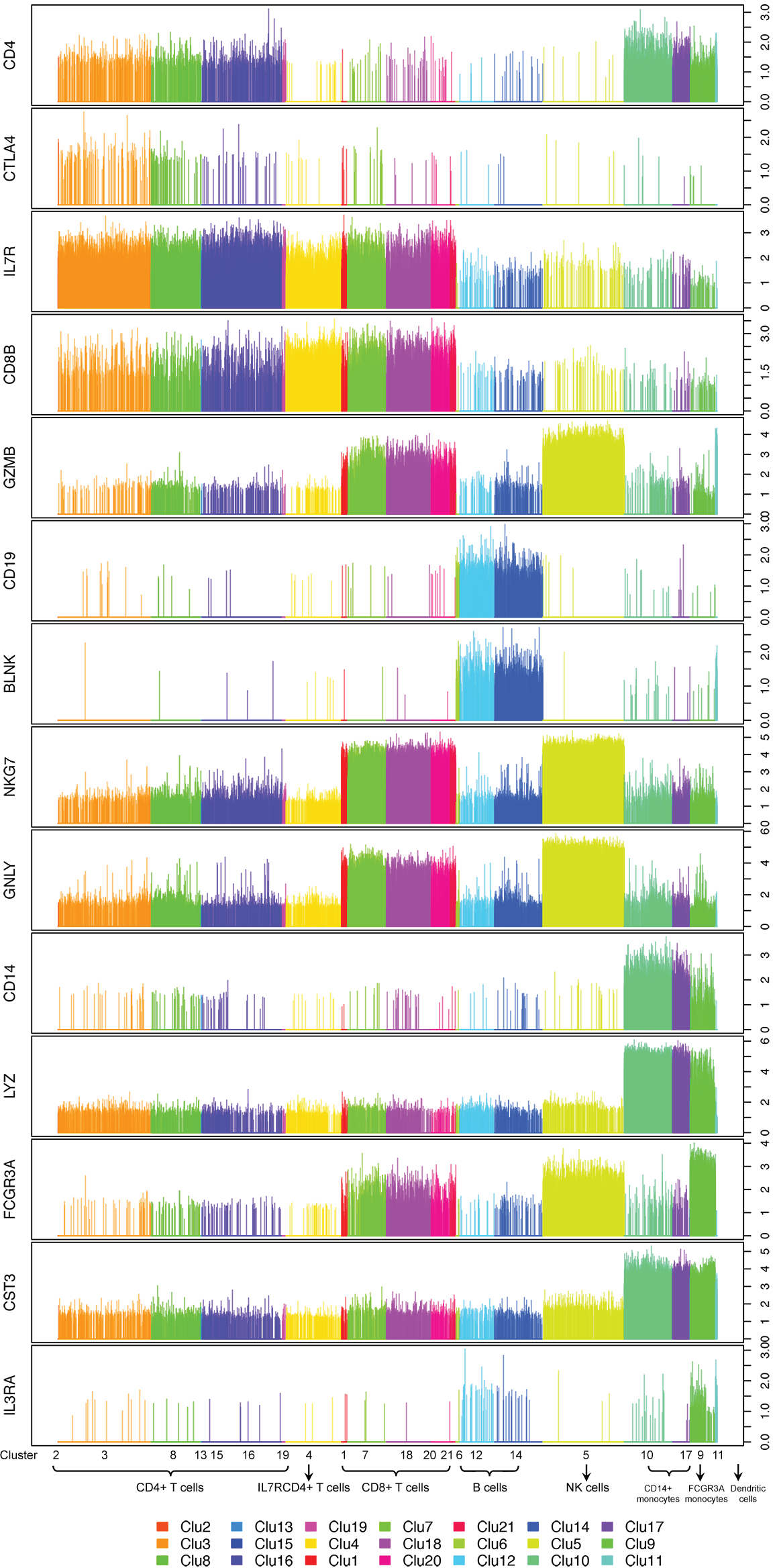
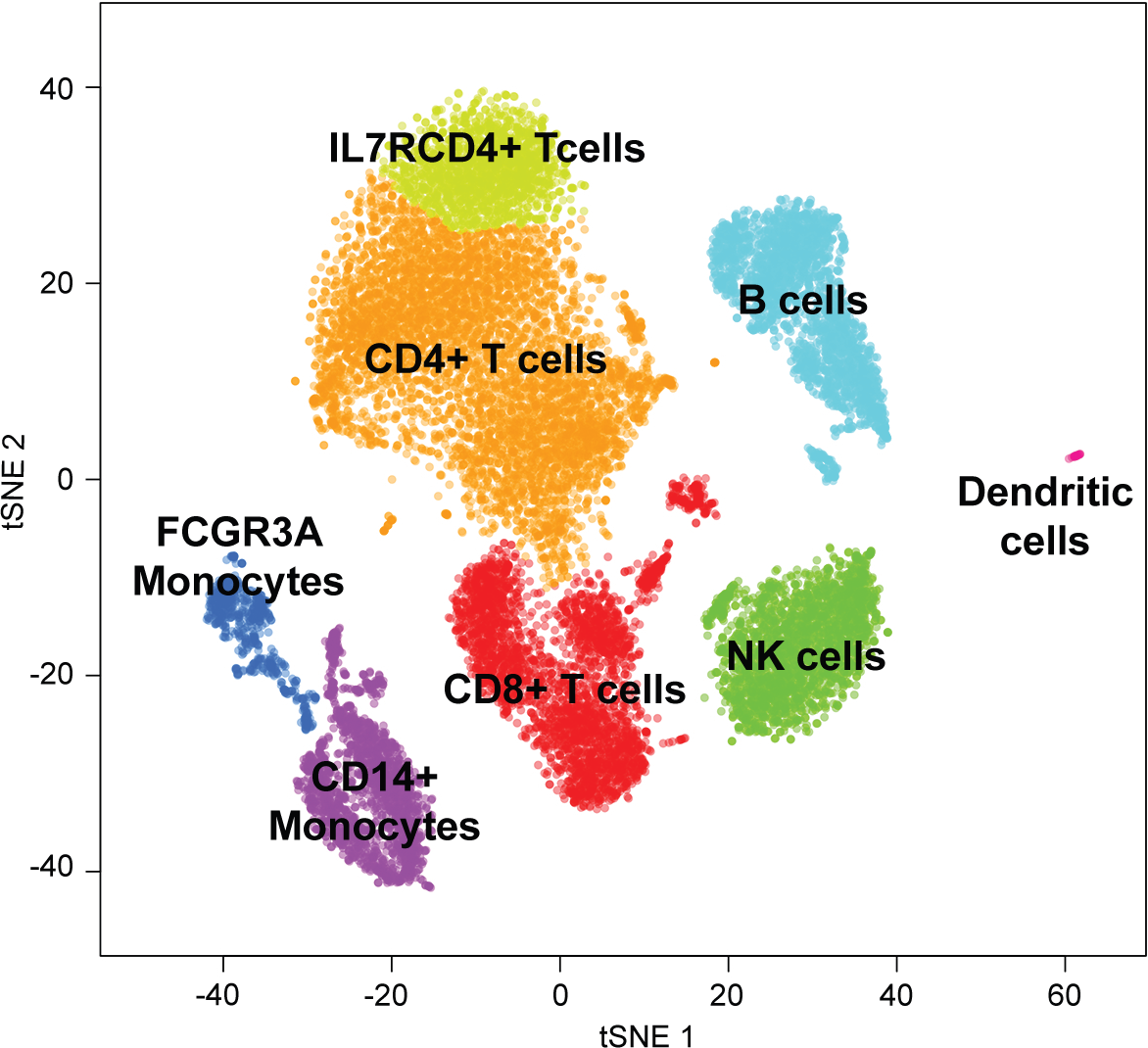
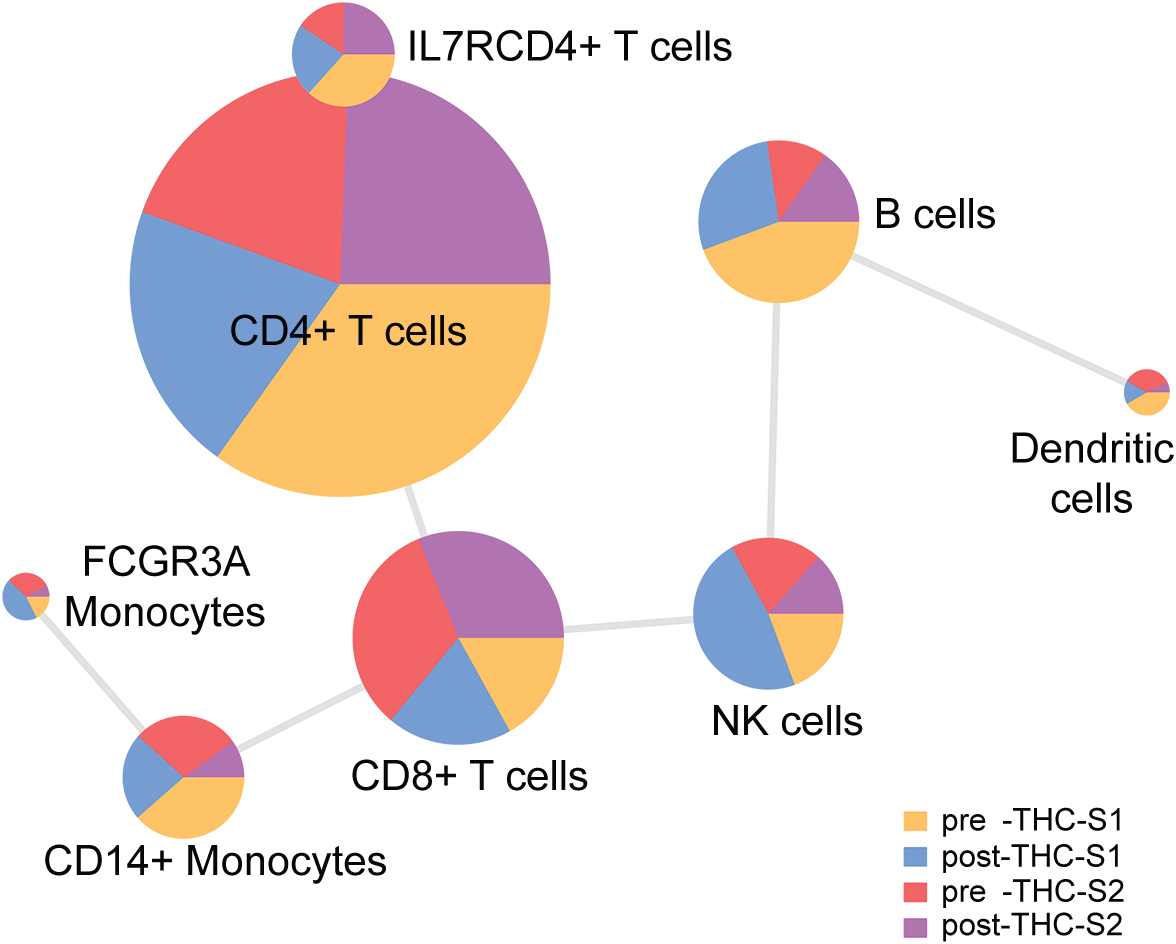
A flow chart illustrating the study design and data analysis strategy. a. Two participants were infused with delta-9-tetrahydrocannabinol (THC). Blood samples were drawn before and 70 minutes after THC infusion. Peripheral blood mononuclear cells (PBMC) were isolated from each sample and 5000 cells subjected to single cell RNA-seq using the 10X Genomics platform. b. tSNE plot showing cell transcriptomic clusters of 15,973 PBMCs in four samples pre- and post-THC infusion: pre-THC-S1; 65: post-THC-S1; pre-THC-S2; and post-THC-S2. Cell number in each sample are presented. The plot indicates a batch effect by participants showing that cell clustered by two participants. c. After removal of batch effects from two participants, 15,973 cells clustered into 21 groups by single cell transcriptomic profile. d. Examples of differentially expressed marker genes in each cell cluster. The clusters with similar marker gene profiles in a given cell type were assigned to the same cell type: CD4+ T-cells Clu(2,3,8,13,15,16,19); IL7RCD4+ T-cells Clu(4); CD8+ T-cells Clu (1,7,18,20,21); B cells Clu(6,12,14); NK cells Clu(5) CD14+ monocytes Clu(10,17); FCGR3A monocytes Clu(9) DCs Clu(11). e. Cell type identification by generalized linear modeling (GLM) cell mapping approach and tSNE plotting. A panel of reference genes for each cell type were selected from previously published studies. GLM tested the association of cell type and marker gene expression in each cell. Significance association is set at p<0.02 and t>0. Each individual cell is assigned to a cell type based on the predominant proportion of marker genes in a given cell type. Small cell clusters are merged into the closest cell type in the tSNE plot. The cell mapping deconvolutes cells to eight PBM cell types: CD4+ T-cells, ILR7+/CD4+ T-cells, CD8+ T-cells, B cells, natural killer cells, CD14+ monocytes, FCGR3A monocytes, and DCs. f. Percentage of cell numbers from each sample in a given cell type: pre-THC-S1, post-THC-S1, pre-THC-S2, post-THC-S2.

Cells (n=15,973) clustered by participant, not by experimental condition (**Figure 1b**), indicating that transcriptomic variability between individuals is greater than variability introduced by a single THC dose. We then removed batch effects using Seurat ^23^ and surrogate variable analysis ^24^ methods and all 15,973 cells clustered into 21 groups (**Figure 1c**, **Figure S1**). To assign cell clusters to cell types, we used a generalized linear model (GLM)-based cell mapping approach with cell-type “marker” genes curated from the literature (see Methods). Briefly, we selected a reference gene panel based on known cell type-specific gene profiles ^22, 25^, then used GLM to test the association of gene expression in each cell with the known marker genes(Figure S2, Table S1). Each cluster was assigned a cell type based on the highest percentage of significant cells (**Table S2**). Expression of marker genes differed significantly in cell types (**Figures 1d**, **S2**). This approach unambiguously deconvoluted the 15,973 cells among 21 clusters into eight cell subtypes: CD4+ T-cells (34.6%), IL7RCD4+ T-cells (8.4%), CD8+ T-cells (17.4%), B cells (13.2%), natural killer (NK) cells (12.3%), CD14+ monocytes (10.0%), FCGR3A monocytes (3.9%), and dendritic cells (DC) (0.3%) (**Figure 1e**). The proportions of each cell type among the participant samples pre- and post-THC infusion are presented in **Figure 1f** and **Table S3**. This robust cell type identification allowed us to examine THC-regulated gene expression in each cell type.

We next applied LMM to detect individual genes affected by THC infusion, with participant included as a random variable to limit confounding effects from an individual’s genomic background. We identified 294 transcriptome-wide significant genes in eight cell types changed by THC infusion (false discovery rate, FDR<0.05) (**Figure 2a**). DCs and FCGR3A monocytes were excluded from further analyses because no gene reached transcriptome-wide significance in DCs and both cell types had low frequencies. Among the 294 DEGs, 69 were observed in at least two cell types while 225 were significant in unique cell types (**Figure 2b**; **Table S4-S11**). Overall, THC infusion resulted in more upregulated genes than downregulated genes.

**Figure 2.**
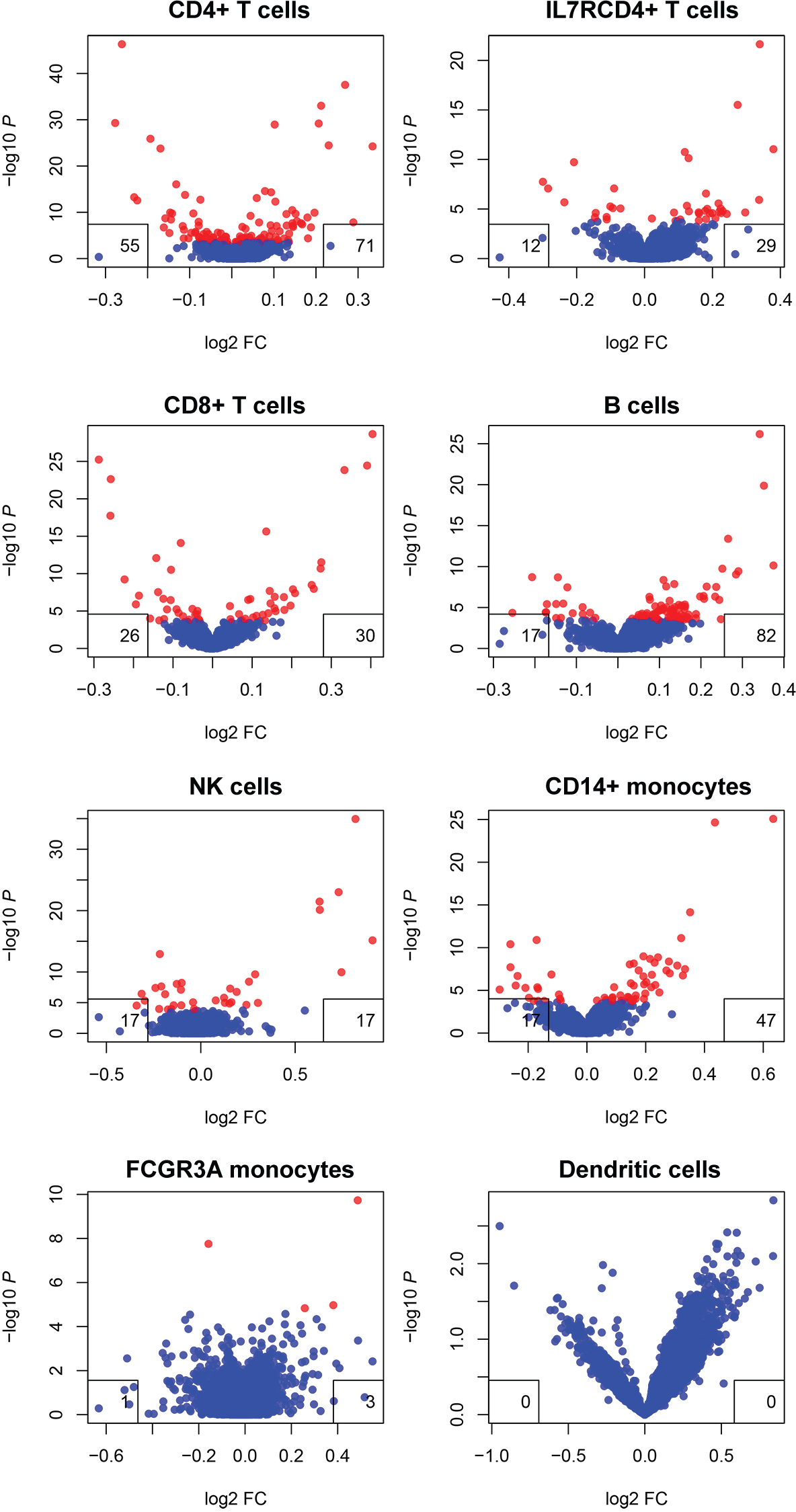
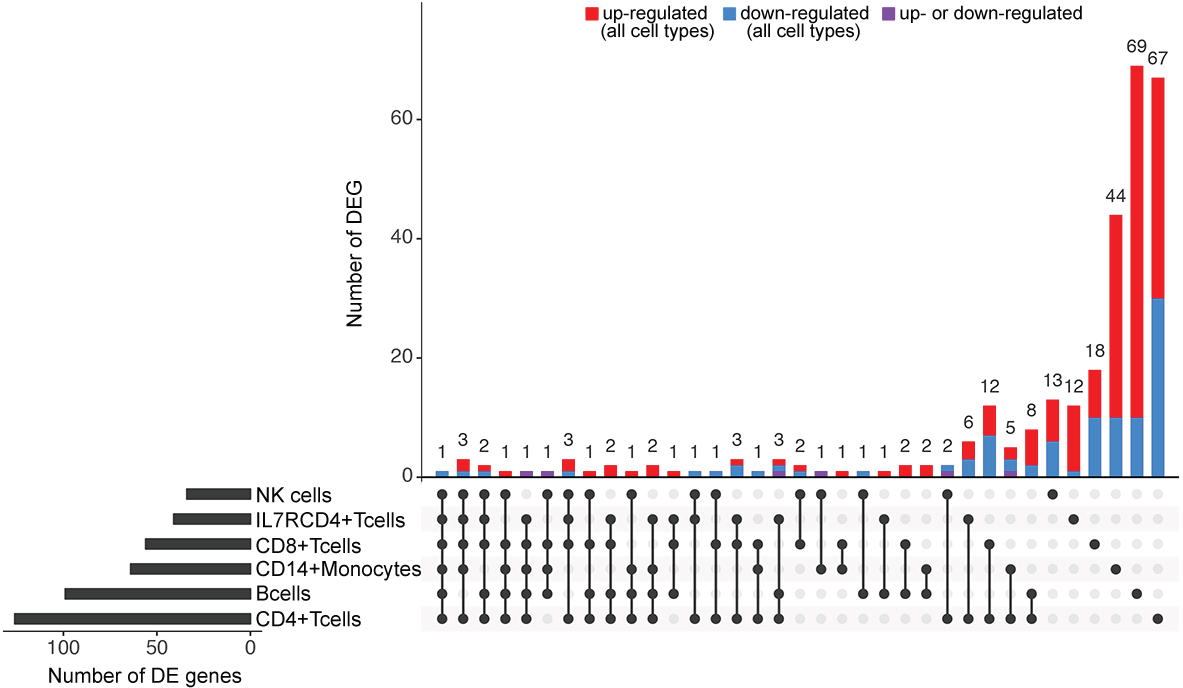
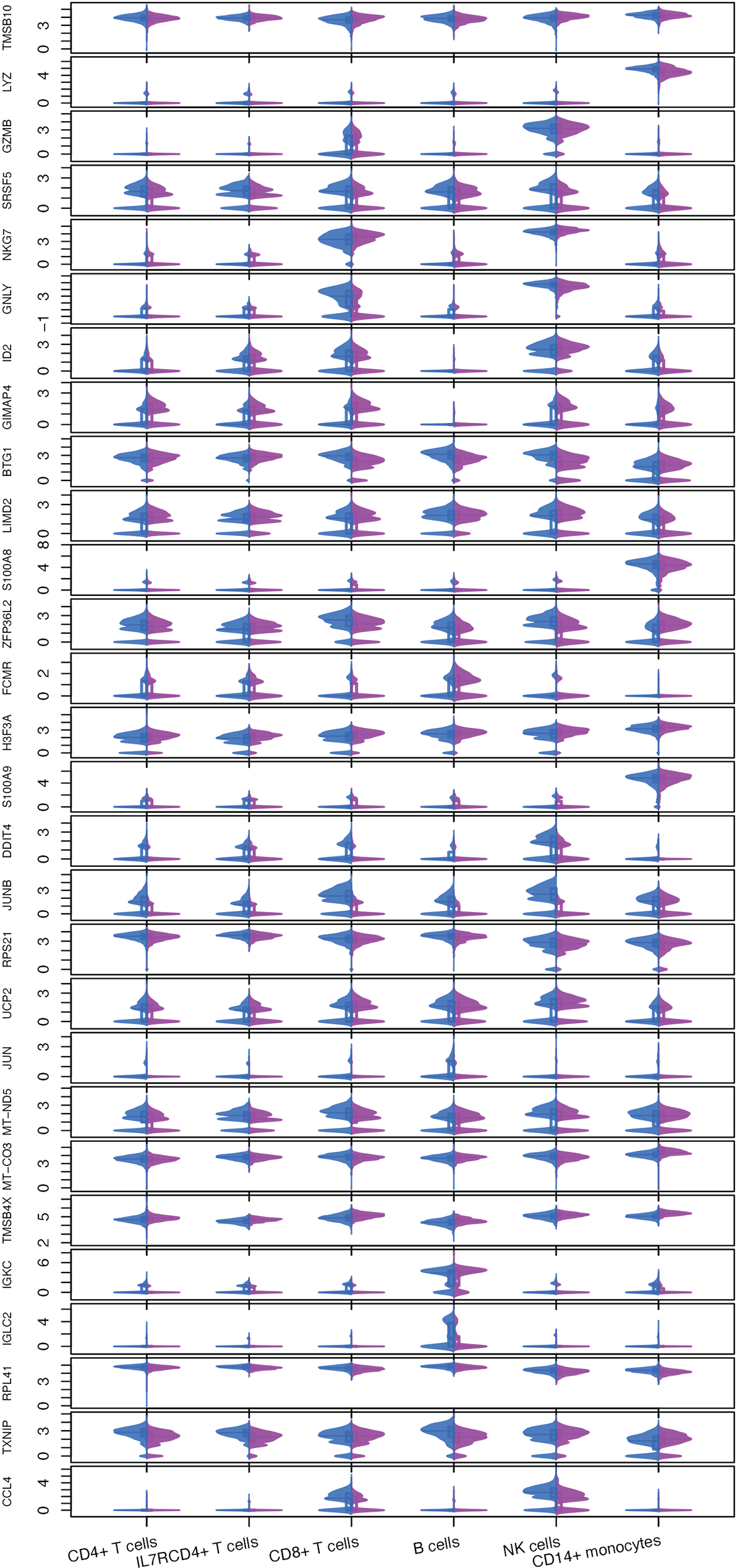
Single cell transcriptome profiling reveal gene expression affected by a single dose of delta-9-tetrahydrocannabinol. a. Linear Mixed Regression analysis identified differential expression of genes affected by THC infusion in eight cell types of peripheral blood mononuclear cells: CD4+ T-cells, ILR7CD4+ T-cells, CD8+ T-cells, B cells, natural killer cells, CD14+ monocytes, FCGR3A monocytes, and DCs. Differentially expressed genes were identified by applying linear mixed modelling (false discovery rate, FDR<0.05). Each inset box denotes the number of up-regulated (right box) and down-regulated (left box) genes. No differentially expressed genes were found in DCs. b. Number of differential expressed genes (DEGs) in six major cell types. X-axis represents DEGs in each cell type; Y-axis represents number of DEG. A total of 69 DEGs are shared in at least two cell types, while 225 DEGs are unique for individual cell type (FDR < 0.05). Among shared DEGs in multiple cell types, up(red)- or down(blue)-regulated genes by THC are in the same direction across cell types except three DEGs (purple). c. Violin plots showing common differentially expressed genes between pre-THC and post-THC in at least three cell types. X-axis represents cell type; Y-axis represents expression level for each gene. Blue: pre-THC samples; pink: post-THC samples.

We sought to identify THC-regulated genes common across the six common cell types. We found 28 DEG in at least three cell types (**Figure 2c**). The majority of the DEG showed consistent directions of regulation by THC in different cell types; only three genes displayed opposing regulation in CD14+ monocytes compared to other cell types (*TMSB4X*, *JUNB, TXNIP*). A group of THC-regulated genes have functions in the domains of immune response and inflammatory process. For example, expression of *S100A9* and *S100A8*, which play prominent roles in the regulation of proinflammatory processes and immune response^26–28^, decreased in response to THC infusion in five cell types. A major HIV-1 suppressive gene, *CCL4*, displayed increased expression after THC infusion. THC decreased expression of *GNLY* that is involved in activating antigen presentation.^29^ The altered expression of genes involved in humoral immunity were also observed (*IGLC2, IGKC)*. Changes in expression of these genes suggest that THC activates the adaptive immune system shortly after administration, in a line with findings showing an immunomodulatory effect that is more complicated than solely immunosuppression.^3, 8, 30–33^

Among the 28 DEG shared in at least three cell types, a subset supports previous findings that acute THC exposure inhibits cell proliferation and induces apoptosis. Genes responsible for cell death were upregulated (*BTG1*,^34^ *DDIT4*,^35^ *GZMB*) and genes involving in cell growth and differentiation were down-regulated (*TMSB10*^36^, *RPS21*,^37^ *RPL41*^38^ by THC exposure. The alteration of these genes in distinct cell types may indicate potentially deleterious effects of THC on cell differentiation and survival.

Given that the majority of THC-regulated DEG are unique to each cell type, we then focused on DEG and co-expression networks in each cell type. DEG were determined both in individual clusters in each cell type and among all cells in each cell type. We leveraged gene-gene relationships cataloged in the Kyoto Encyclopedia of Genes and Genomes (KEGG) database^39^ and constructed gene networks independently in pre- and post-THC samples in each cell type (FDR<0.05). Hub genes in each cell-type based network were defined as a node (gene) with at least four edges (gene links) in at least one condition (pre-, post-THC, or both). We then performed Gene Ontology (GO) term enrichment analysis for each cell-type based co-expression network (FDR<0.05).

In CD4+ T-cells, significant DEG are involved in cytotoxic T-cell activation (*IL7R*^40^), histone modification (*H3F3B*^41^), and transcriptional regulation (*MYC*^42^). We observed three cell sub-groups (cluster 3, 8, and 16; **Figure 3a**; **Table S12**). Cluster 8 showed a distinct DEG pattern from clusters 3 and 16. Co-expression analysis identified 40 nodes with 33 edges enriched on 14 GO terms including immune response, cell surface receptor signaling pathway, cellular response to stimulus. Two hub genes in the network, *CCR7* (significantly connected with *CCR* [*CCR2, CCR6, CCR10*] and *CXCR* [*CXCR3*, *CXCR4*] family genes) and *HLA-A* (significantly correlated with three other *HLA* genes [*HLA-B, HLA-E, HLA-F*], were significantly affected by THC infusion (**Figure 3b**), showing that acute THC exposure perturbed gene-gene relationships in CD4+ T-cells and appears to increase the gene connectivity involving cell-mediated immunity.

**Figure 3.**
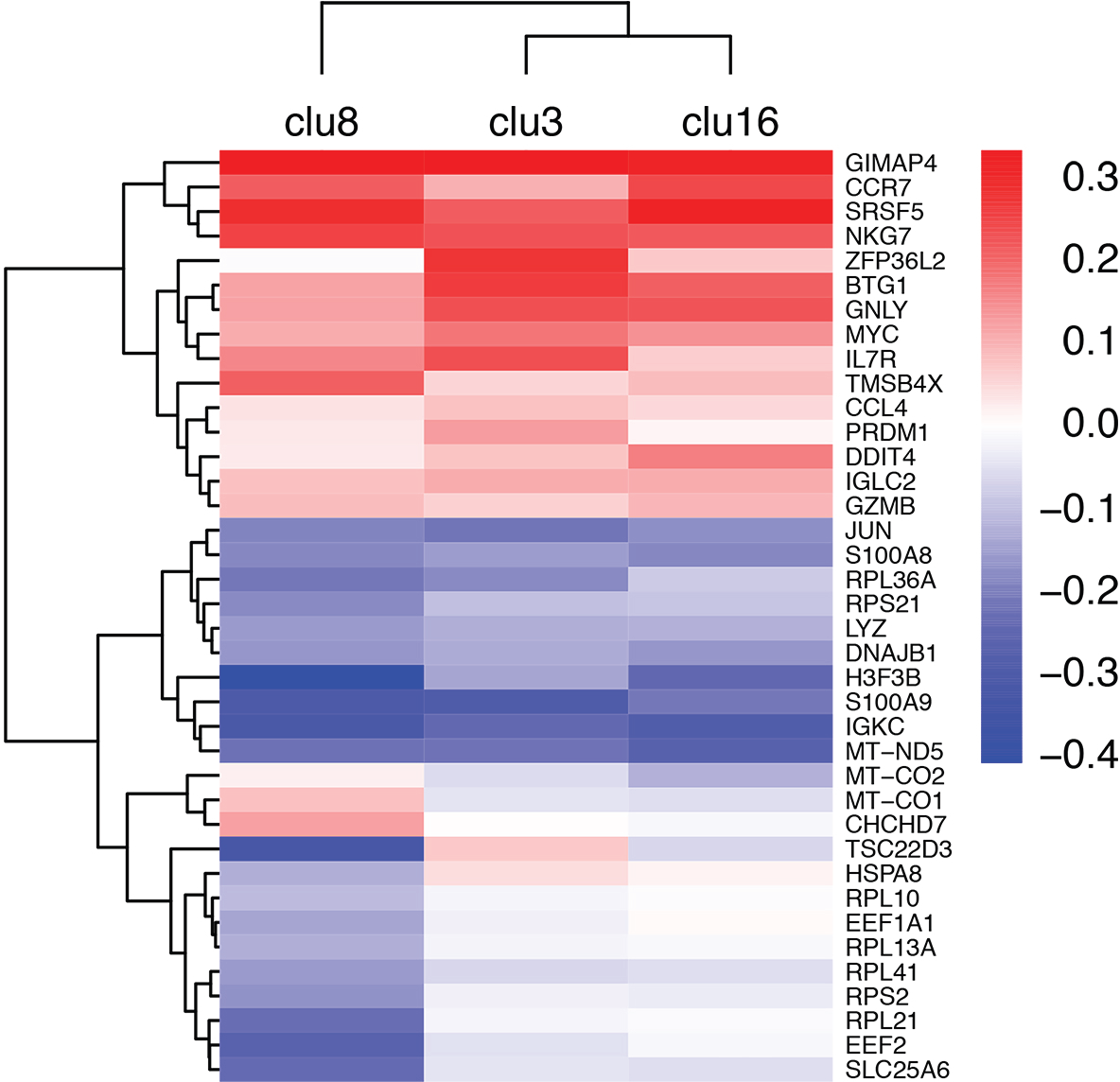
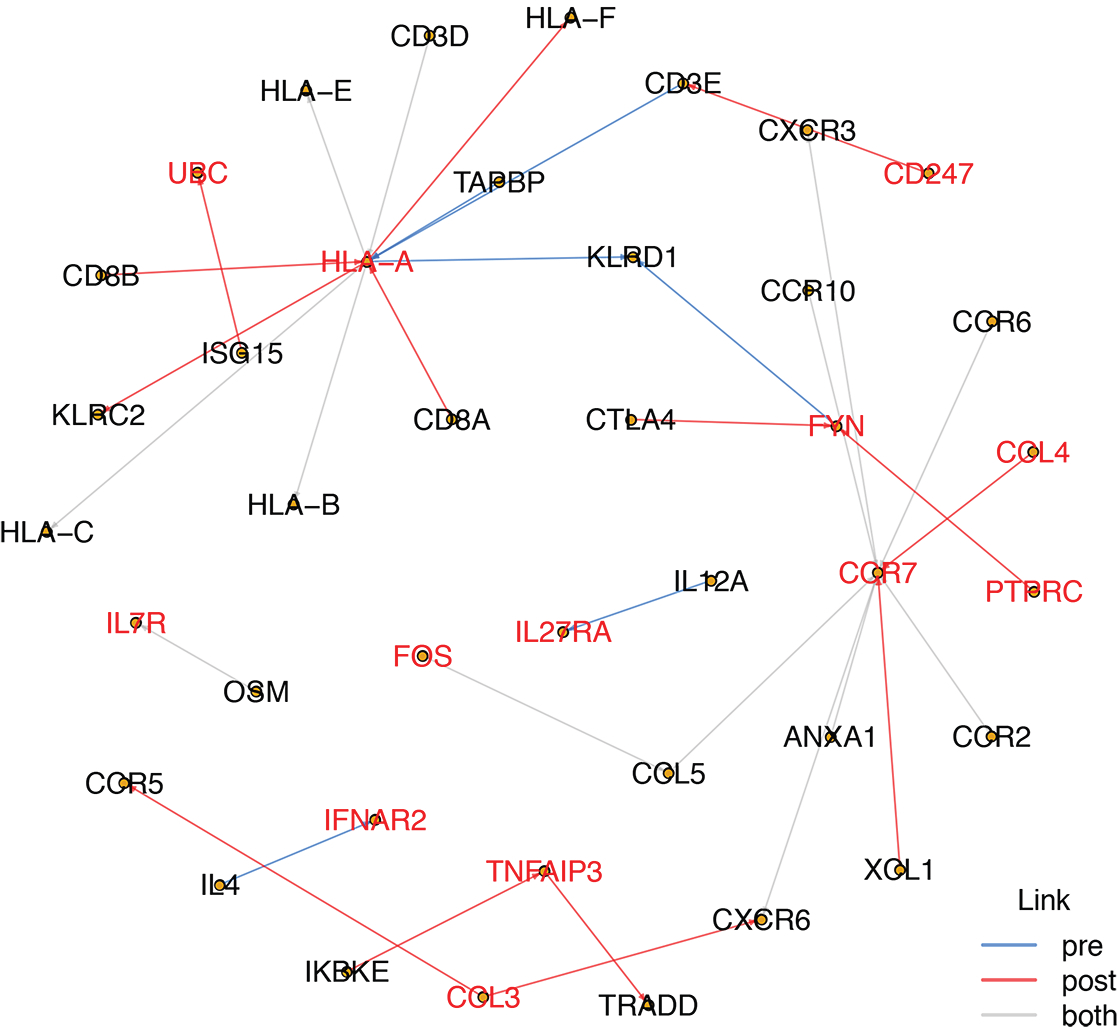
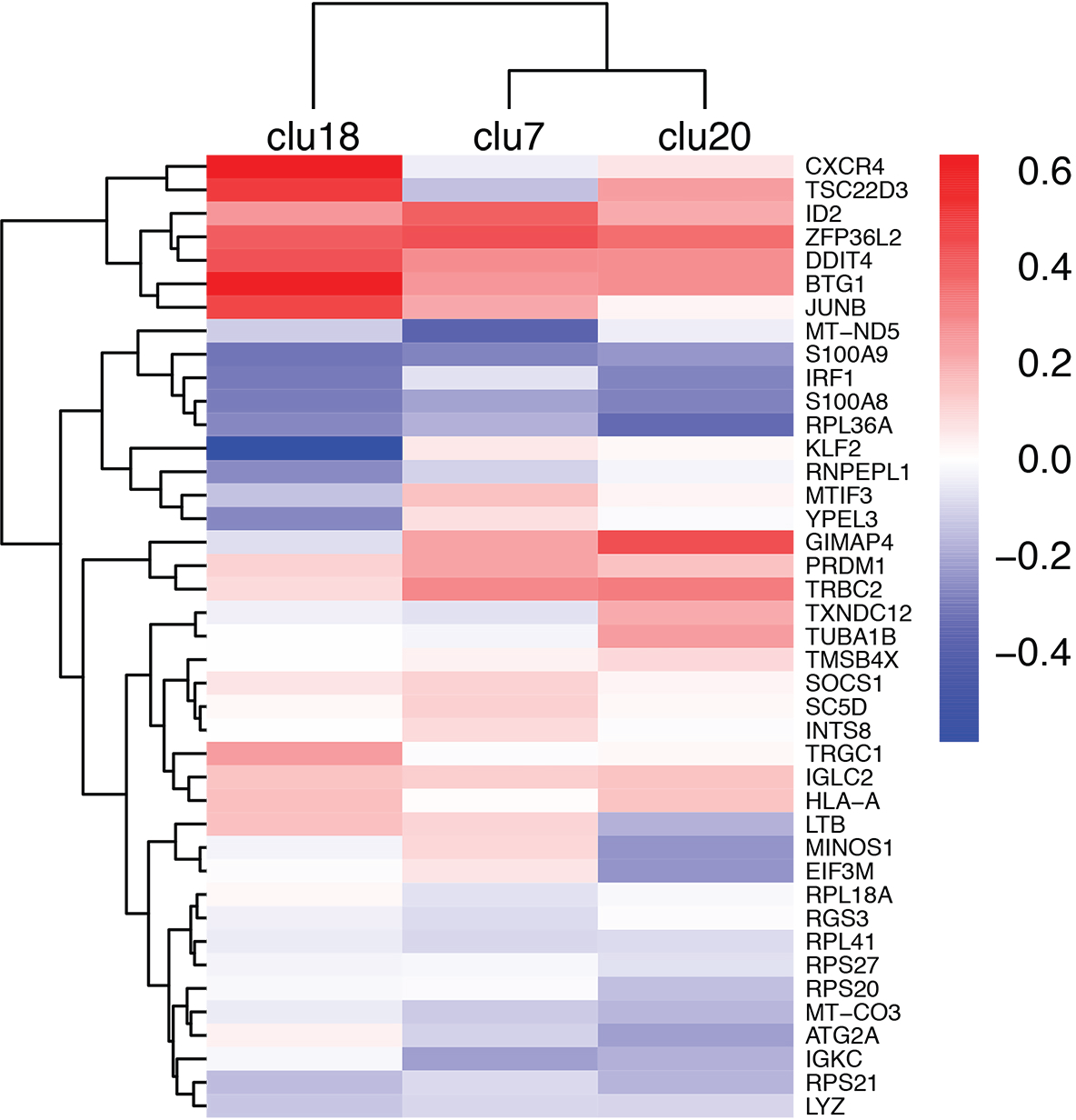
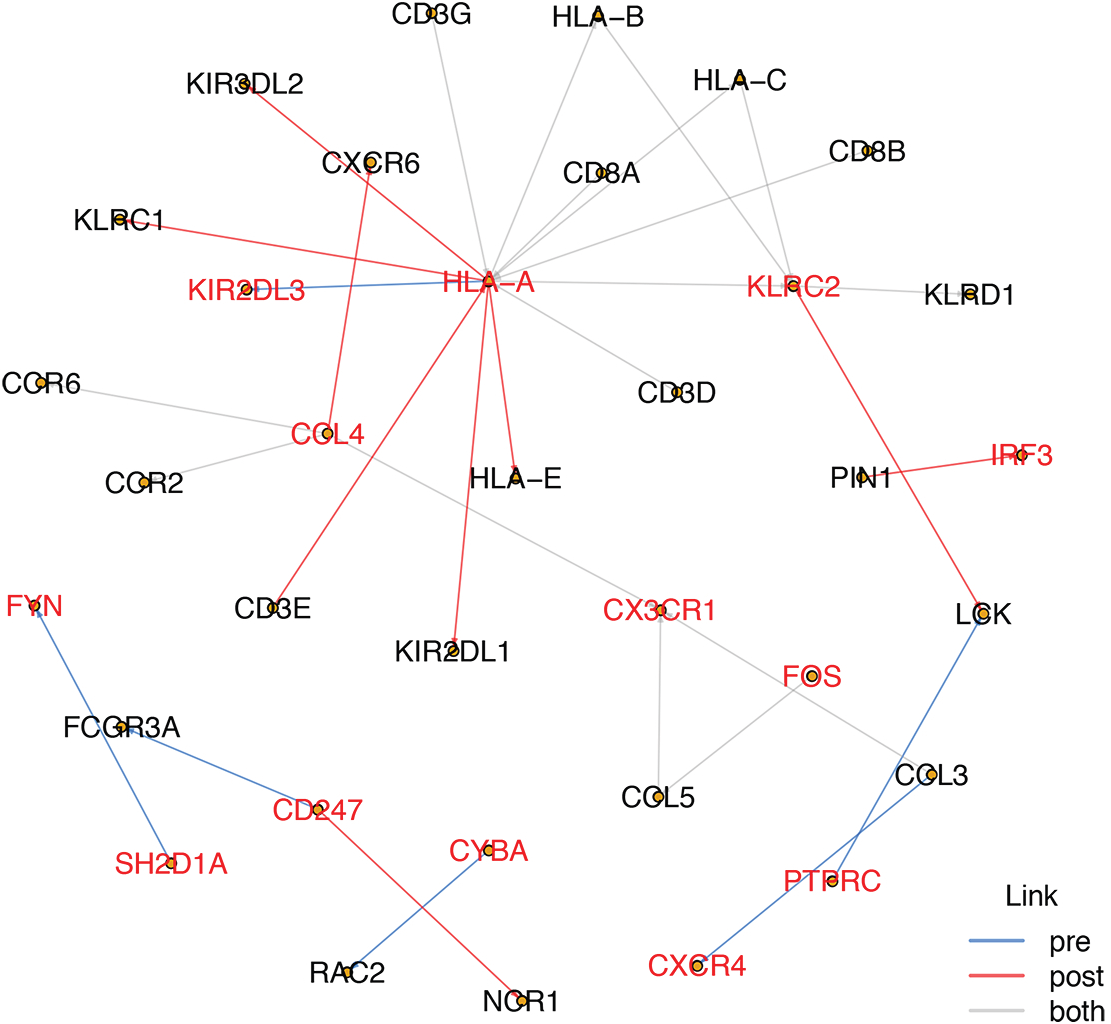
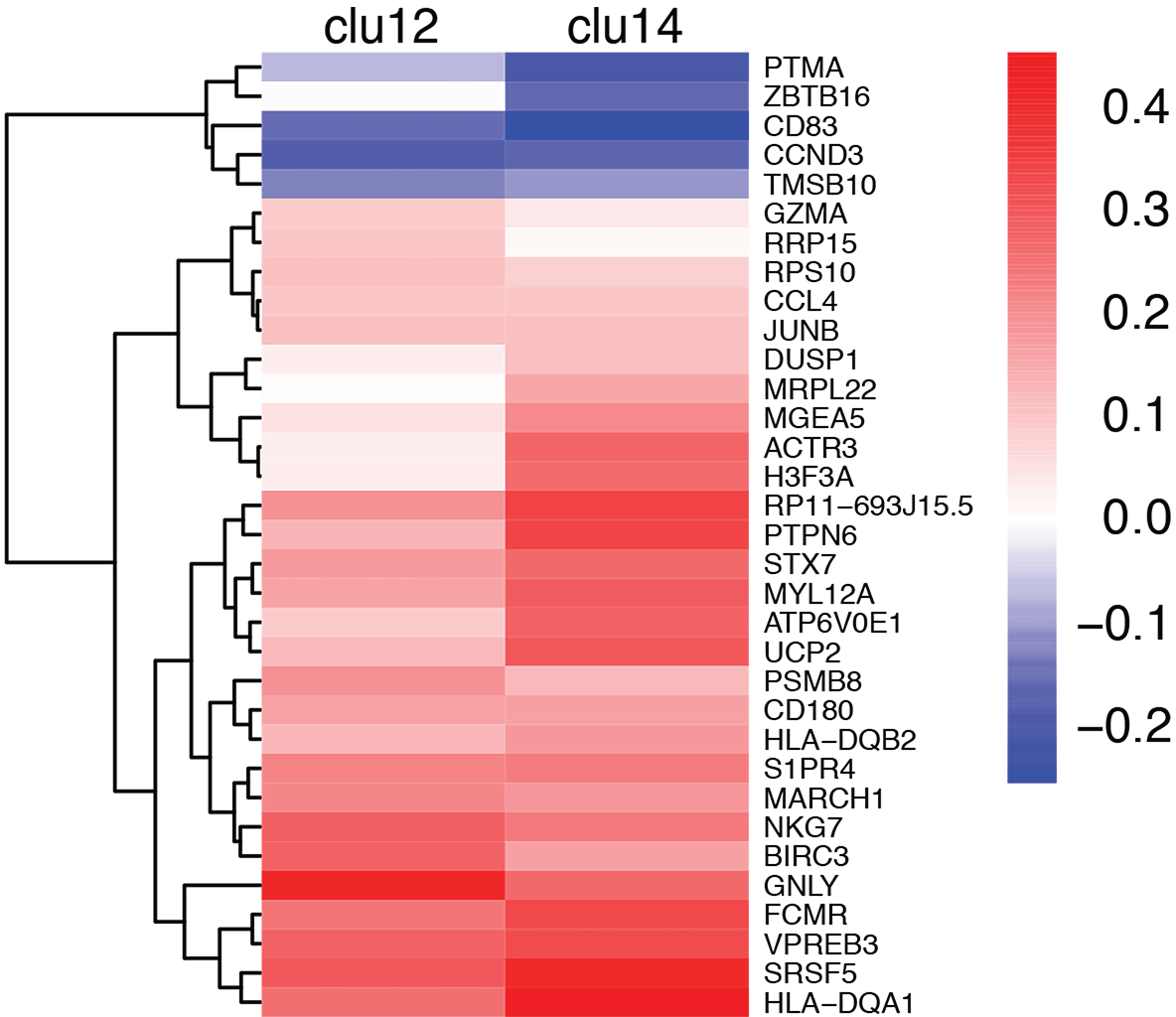
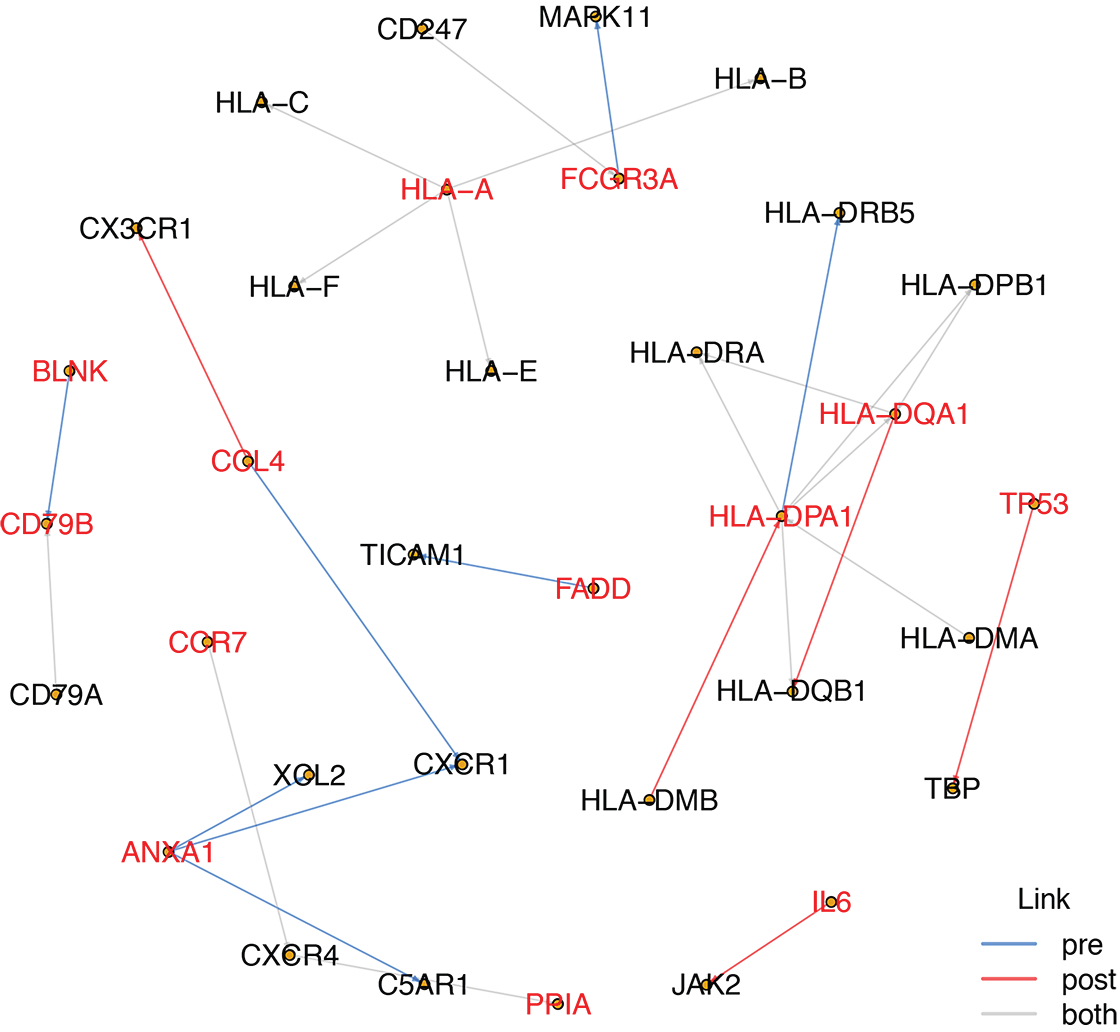
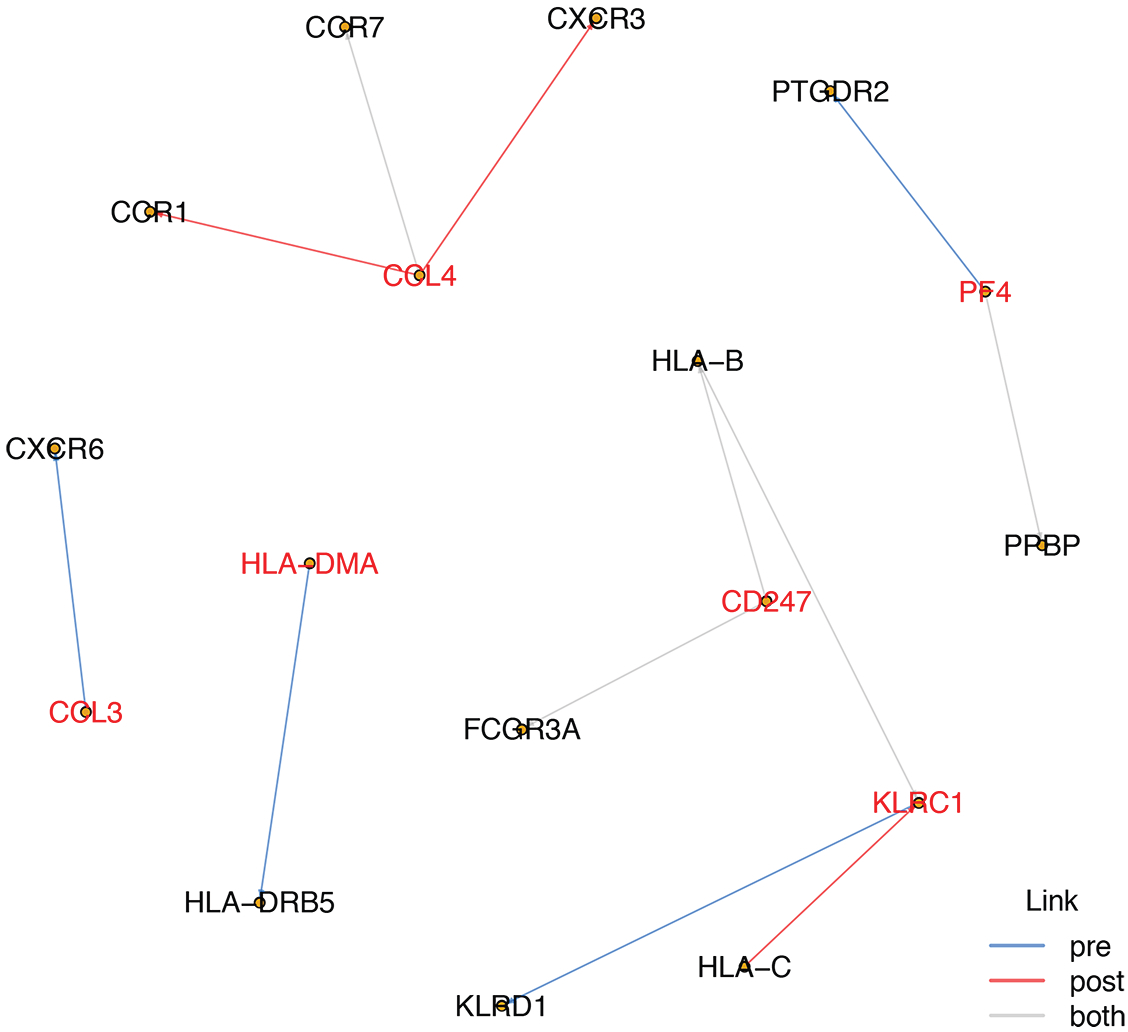
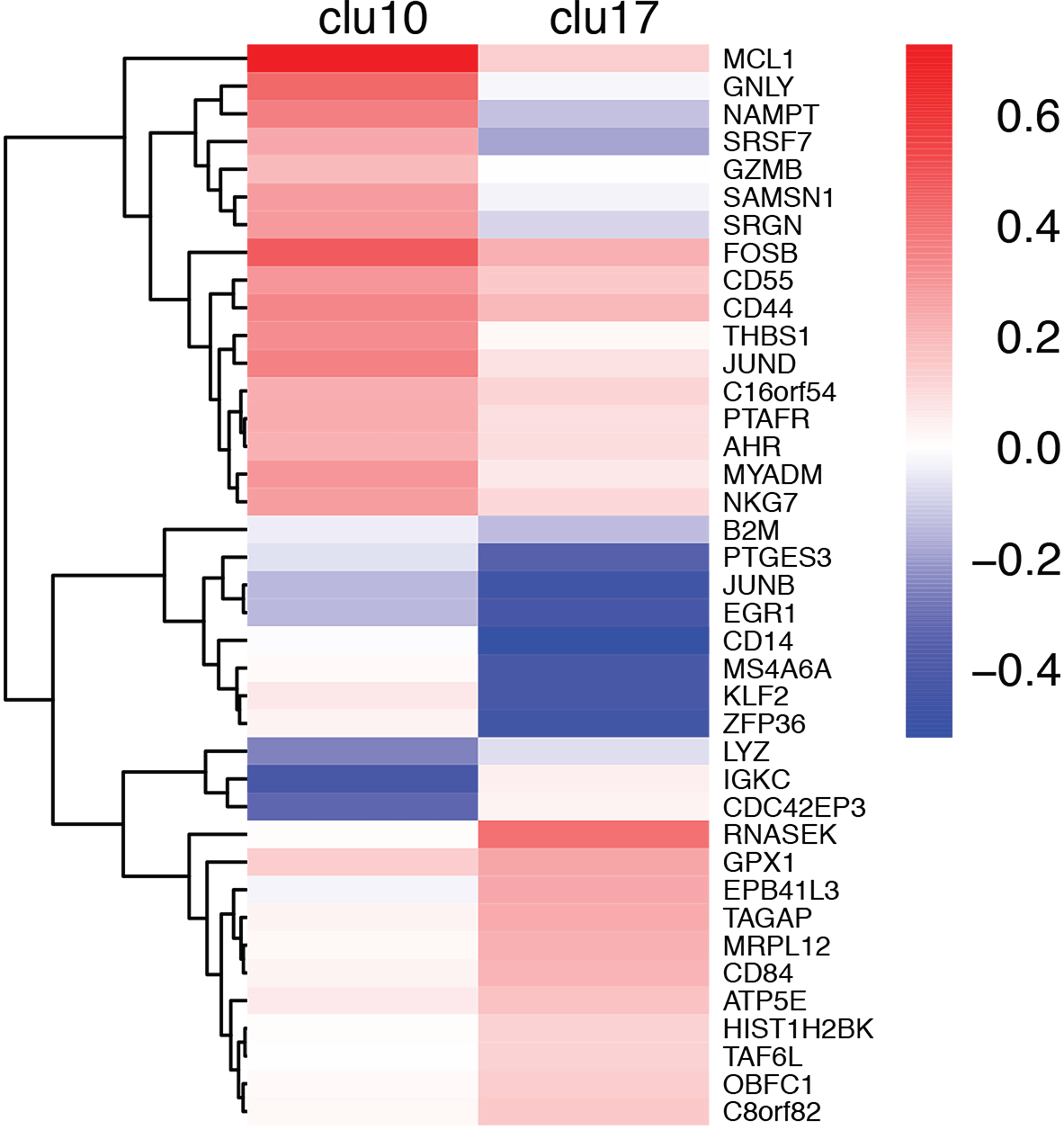
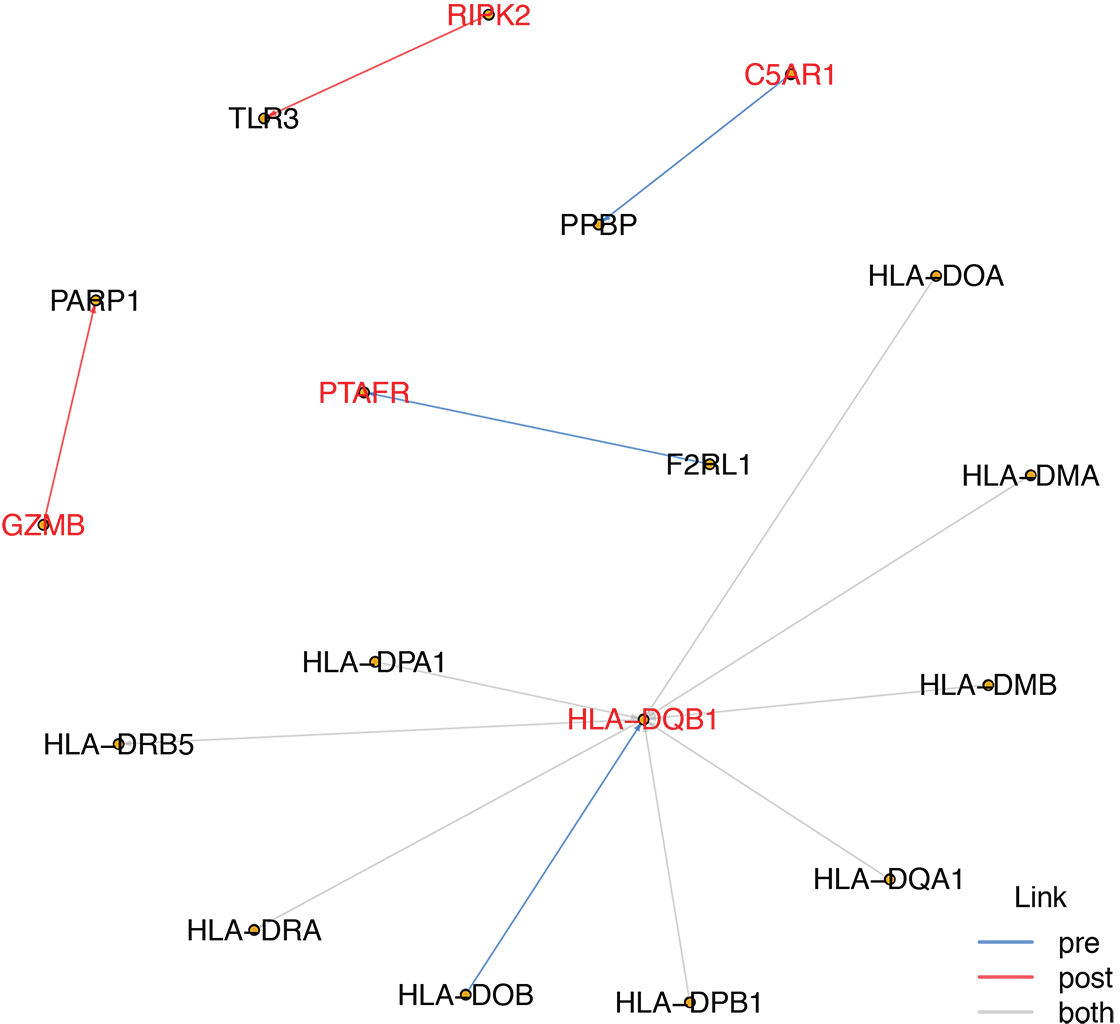
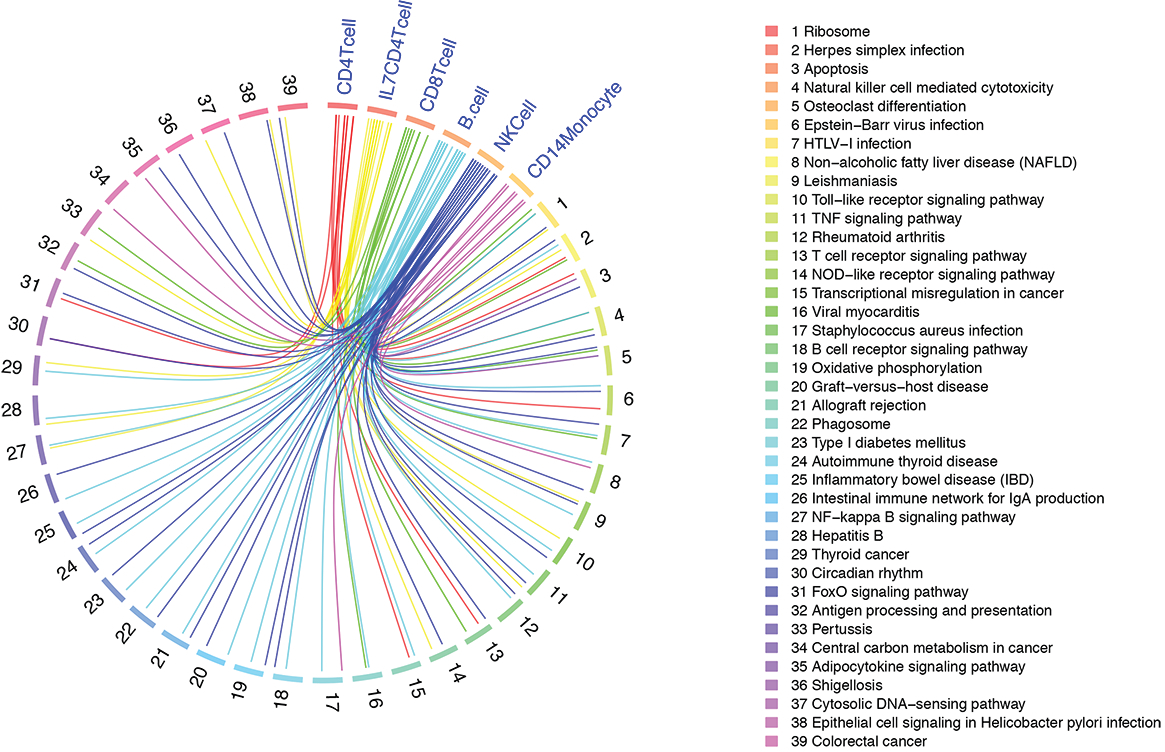
Cell-type based differential gene expression, gene-gene correlation, and biological pathways influenced delta-9-tetrahydrocannabinol. a. Heat map showing three subtypes of CD4+ T-cells affected by THC. Color bar represents log2 fold change between post-THC and pre-THC samples. Each row represents significant genes for top 20 genes in each cell cluster. Cluster 8 shows a distinct pattern of differential gene expression from those observed in clusters 3 and 6. In cluster 8, nine genes encoding ribosomal protein are decreased in expression by THC. b. Co-expression network in CD4+ T-cells. Genes with nominal p<0.001 from linear mixed regression model are selected to construct a co-expression network. Gene-gene links were derived from KEGG: Kyoto Encyclopedia of Genes and Genomes – GenomeNet. Significance is set at false discovery rate (FDR)<0.05. A total of 40 nodes and 33 edges were observed. Two hub genes, *CCR7* and *HLA-A*, are differentially expressed genes influenced by THC. c. Heat map showing three subgroups of CD8+ T-cells affected by THC. Cluster 20 shows distinct differential gene patterns from clusters 7 and 18. d. Co-expression network in CD8+ T-cells. Genes with nominal p<0.001 in CD8+ T-cells are selected to construct a co-expression network. In CD8+ T-cells, we observed 35 nodes and 31 edges. One hub gene, *HLA-A*, is differentially expressed by THC. e. Heat map showing two subtypes of B cells affected by THC. Each row represents significant differentially expressed genes in B cells. Cluster 12 shows a distinct differential gene expression pattern from that of cluster 14. f. Co-expression network in B cells. Genes with nominal p<0.001 in B cells are selected to construct a co-expression network. A total of 34 nodes and 28 edges are observed. Three hub genes, *HLA-A, HLA-DQA1*, and *HLA-DPA1* are identified. g. Co-expression network in natural killer (NK) cells. We observe 17 nodes and 12 edges. One hub gene, *CCL4*, is significantly differentially expressed following THC infusion. h. Heat map showing two distinct subtypes of CD14+ monocytes (cluster 10 and cluster 17) affected by THC. Color bar represents log2 fold change between pre-THC and post-THC samples. Each row represents significant genes in CD14+ monocytes. i. Co-expression network in CD14+ monocytes. Genes with nominal p < 0.001 in CD4+ T-cells are selected to construct a co-expression network. We observed 18 nodes and 13 edges. One hub gene, *HLA-DQB1*, was identified in both pre-THC and post-THC samples. j. Circus plot showing gene set enrichment analysis using KEGG annotations reveal 39 biological pathways shared in at least 2 cell types. KEGG: Kyoto Encyclopedia of Genes and Genomes – GenomeNet. Significant pathway is declared at false discovery rate (FDR)< 0.05. The legend is the name of each pathway corresponding to each spoke of the circular plot.

In CD8+ T-cells, we identified 18 unique DEGs that involved in immune response and inflammation in response to THC (e.g. *IL32*,^43^ *SOCS1*,^44^ and *IRF1*^45^). Of note, THC infusion resulted in the differential expression of *CXCR4, TSC22D3, DDIT4, BTG1, JUN* in cluster 18 of CD8+ T-cells (**Figure 3c**), suggesting that cells in cluster 18 may function differently in response to THC as compared to the other CD8+ T-cell sub-group. The gene network included 35 nodes that enriched on 12 GO terms (e.g., immune system process), similar to the CD4+ T-cell network. One hub gene, *HLA-A* (strongly correlated with *HLA-B* and *HLA-C* in pre- and post-THC samples), was observed in the network (**Figure 3d**; **Table S13**).

In B cells, genes unique to B cells were observed that are involved in B cell maturation (*VPREB3*^46^), MHC function (*HLA-DQA1*, *HLA-DQA2*), calcium signaling (*CALM2*),^47^ Toll-like receptors (*CD180*),^48^ and response to environmental stress via activating MAP kinase MAPK1/ERK2 (*DUSP1*).^49^ Consistent with *in vitro* findings, we found that acute THC exposure reduced expression of *BCL2* in B cells.^40^ The majority of DEG originated from cluster 14 (**Figure 3e**; **Table S14**), with only 8 DEGs from cluster 12, which were all upregulated by THC. In the co-expression network, we found 34 nodes with 28 edges (**Figure 3f**) enriched on GO pathways including MHC protein complex, immune response, peptide antigen binding. Four hub genes were identified that differed in response to THC (*HLA-A and HLA-BQA1) HLA-DPA1* [became more strongly correlated with *HLA-DQB1* post-THC]*, ANX1* [connected to *CXCL1*, *CXCR4*, *XOL2* in pre-but not post-THC samples]). Notably, two B cell marker genes (*BLNK*, *CD79B*) were correlated in post- but not pre-THC samples.

In NK cells, which is defined by one cluster, many DEGs (e.g. *DDIT4, CCL4, BTG1 ID2*) are involved in immune response and cell proliferation. One DEG unique to NK cells, *CD53*, was downregulated by THC. Genes in the NK cell network (17 nodes and 12 edges; **Figure 3g**) were enriched on 17 pathways including chemokine-mediated signaling pathway, inflammatory response, and chemokine receptor activity. Notably, *KLRC1* regulates specific humoral and cell-mediated immunity and is implicated in the recognition of the MHC class I HLA molecules in NK cells and was correlated with *HLA-C*, which increased in post-THC samples. No hub genes were observed.

In CD14+ monocytes, which were composed of two subgroups (cluster 10 and 17; **Figure 3h**; **Table S15**), THC infusion resulted in unique 44 DEGs, including genes regulating cell fate (e.g. *MCL1*, *FOSB*, *MYADM*). Genes in the CD14+ monocyte network (18 nodes and 13 edges; **Figure 3i**) were enriched on pathways including MHC class II protein complex, antigen processing, immune response, and cellular response to interferon-gamma. One hub gene, *HLA-DQB1*, was observed.

These observations from individual gene and co-expression networks in different cell types suggest a diverse functional response to acute THC exposure in heterogenous immune cells. Significant pathways for all co-expression networks in each major cell type are presented in Table **S16-S21**.

We subsequently performed a cell type-based gene set enrichment analysis (GSEA) using DEGs. We found 39 significant KEGG pathways in at least two cell types (**Figure 3j**; **Table S22-S29**); significant pathways were involved in the domains of immune response, inflammation, and cell survival and apoptosis. Several pathways associated with autoimmune disease (e.g., rheumatoid arthritis) were significant in multiple cell types. The ribosomal pathway, which plays a role cell growth and cellular response to stress, was the most significantly enriched pathway across all five cell types. These results further support the effects of THC on functional domains in immune regulation that have also been associated with immunological disease, although the causality of these relationships is unknown.

Finally, we were interested in understanding how cannabinoid receptor genes co-expressed with other genes in each cell type following THC administration. As expected, *CNR2*, encoding for CBR2, was highly expressed in B cells, followed by NK cells, then CD8+ T-cells, and lowest in CD4+ T-cells (**Figure 4a**).^50^ Little *CNR1* expression was detected in any of the cell types. *GPR55* (cannabinoid receptor 3^51^) showed the highest expression in CD4+ T-cells. We observed gene co-expression between *CNR2* and 84 and 74 genes in pre- and post-THC samples, respectively (**Table S30-S31**). The co-expressed genes in post-THC B cells were enriched on functional domains of immune processes, biological regulation, cell proliferation, signaling, and response to stimulus (**Figure 4b**).

**Figure 4.**
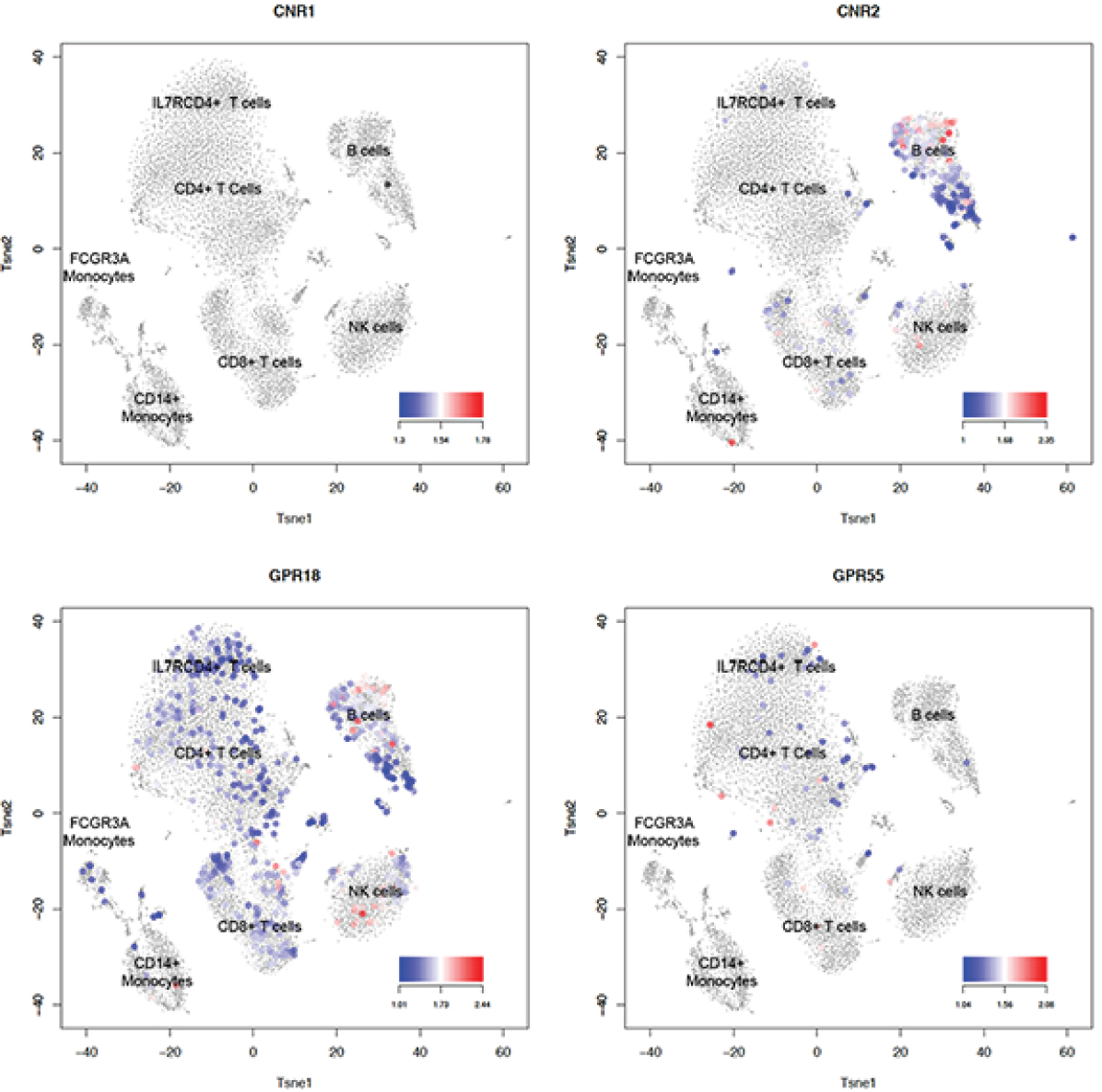
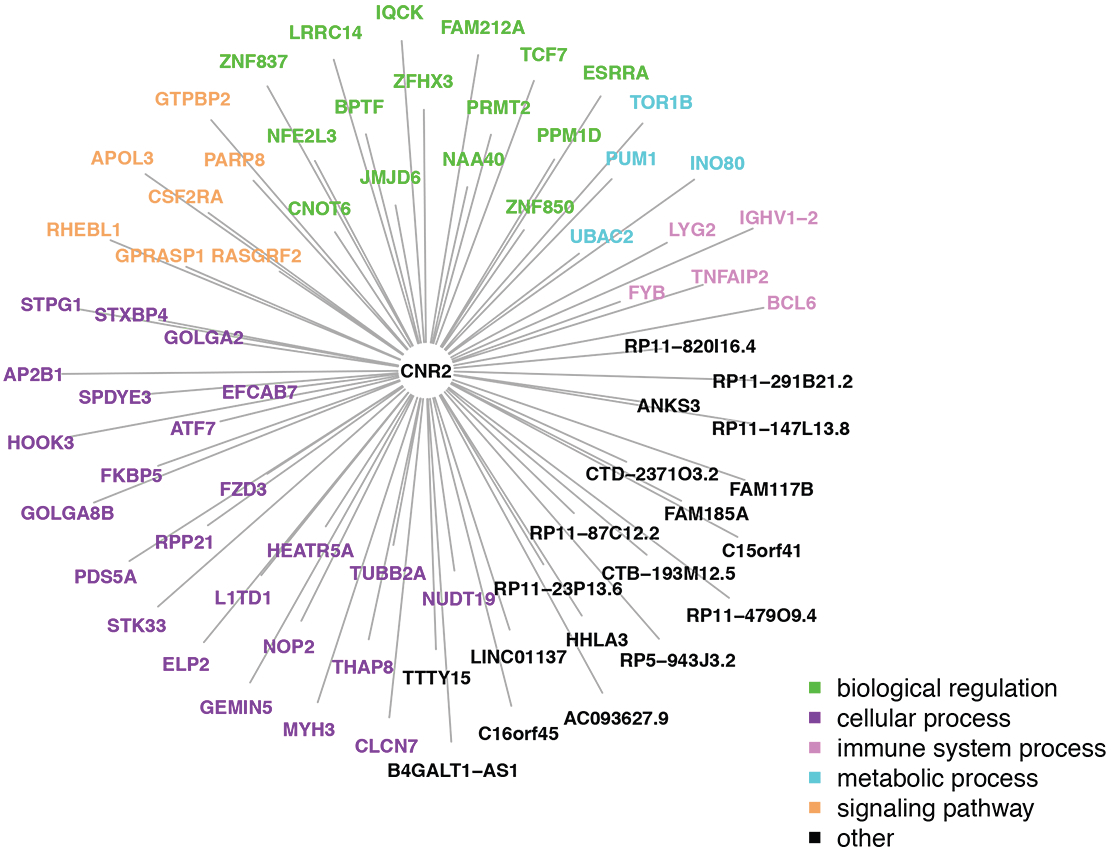
THC receptor gene expression and co-expression network in B cells. a. tSNE plot showing THC receptor gene expression in distinct cell types of peripheral blood mononuclear cells. *CNR1* is observed little expression in any cell types. *CNR2* is highly expressed in B cells, followed by nature killer cells, CD8+ T-cells, and minimally expressed in CD4+ T-cells. *GPR55* is highly expressed in CD4+ T-cells more than other cell types. *GPR18* is widely expressed in peripheral blood mononuclear cells. b. *CNR2* expression is significantly correlated with 74 genes in B cells. The genes co-expressed with *CNR2* are involved in functions of biological regulation, cellular process, immune system process, metabolic process, and signal functions.

In summary, our results from a well-controlled, within subject human study demonstrate transcriptomic regulation in distinct immune cells following administration of a single dose of THC. Our study design and analytical approach enabled the identification of common and cell-type specific DEGs regulated by THC. Subjective, behavioral and cognitive tests confirm that the dose of THC was relevant to but limited the confounding effects of other compounds in cannabis. The cell type-specific gene and co-expression patterns revealed by scRNA-seq showed little overlap across cell types, which would have been obscured by bulk RNA-seq analysis. Cell type-based DEGs, co-expression networks, and GSEA revealed important THC effects on immune function, cytokine production, signal transduction, and cell apoptosis and survival. The study provides a foundation for future studies of cell type-specific immunologic effects of cannabis and cannabinoid constituents. Studying the effects of chronic medicinal or recreational cannabinoid exposure or the effects in immune disorders (e.g., HIV) using our approach are also warranted. These findings highlight the complexity of cannabinoid effects on immune function and nuance of immune cell types that may have medical relevance.

## Methods

The study was conducted at the Neurobiological Studies Unit (VA Connecticut Healthcare System, West Haven, CT) with the approval of the Institutional Review Boards at VA Connecticut and Yale University, and the Protocol Review Committee of the Department of Psychiatry, Yale University. The study was amended to include prospective measures addressing safety.

### Participants

Two healthy participants were recruited from the community by advertisements. Both participants were male, 21-year old, and of European American descent. Subjects were informed about the potential for psychosis, anxiety, and panic. After obtaining informed consent, subjects underwent a structured psychiatric interview for DSM-IIIR ^52^ and were carefully screened for any DSM-IV Axis I or Axis II lifetime psychiatric or substance abuse disorder (excluding nicotine) and family history of major Axis I disorder. The history provided by subjects was confirmed by a telephone interview conducted with an individual (spouse or family member) identified by the subject prior to screening. In order to avoid exposing cannabis-naïve individuals to a potentially addictive substance, only subjects who had been exposed to cannabis but did not meet lifetime criteria for a cannabis use disorder were included. Past month cannabis use was quantified using a time-line-follow-back approach. Finally, subjects underwent a general physical and neurologic examination, EKG, and laboratory tests (serum electrolytes, liver function tests, complete blood count with differential and urine toxicology). Subjects were instructed to refrain from caffeinated beverages, alcohol, and illicit drugs from 2 weeks prior to testing until study completion. Urine toxicology was conducted on the morning of each test day to rule out recent illicit drug use.

### Procedure

Subjects received 0.03mg/kg of THC, the principal active ingredient of cannabis.^18, 19^ This dose equivalent to 2.1 mg in a 70 kg individual has been shown in previous studies to produce effects consistent with the known effects of cannabis, in a safe manner.^53–55^ THC was administered on its own, without the >450 other chemical constituents of cannabis because THC is the principal active constituent of cannabis and the other chemical constituents could render the results challenging to interpret. The intravenous route of administration was chosen to reduce inter and intraindividual variability in plasma THC levels with the inhaled route.^56^ Timeline of behavioral assessment and blood draw is presented in **Figure S3**. Subjects were attended to by a research psychiatrist, a research nurse, and a research coordinator. Clear ‘stopping rules’ were determined a priori and rescue medication (lorazepam) was available if necessary. Medical condition and psychiatric status of participants were monitored closely during and after THC challenging. Subjective and clinical ratings were repeatedly assessed.

### Medical and behavioral assessment

Vital signs, Cannabis intoxication, Psychotic symptoms, Perceptual alteration, and Cognitive test battery were measured prior, during, and post THC infusion as illustrated in **Figure S3**.

### Single cell RNA sequencing in 10X Genomics platform

PBMCs from pre-THC (N=2) and post-THC samples (N=2) from fresh whole blood were isolated at the same time using a standard protocol. The cells were washed twice with phosphate-buffered saline containing 0.04% bovine serum albumin, and the final cell concentration was adjusted to 1000 cells/mL for library preparation. 5000 cells/sample were prepared for single cell capture.

#### Sequencing data processing and quality control

We used the 10X Genomics Chromium Single Cell 3’ v2.0 platform to prepare individually barcoded single-cell RNA-Seq libraries following the manufacturer’s protocol. Library size was confirmed with Agilent Bioanalyzer High Sensitivity DNA assay (PN:5067-4626), Invitrogen dsDNA HS qubit assay to evaluate dsDNA quantity (PN: Q32854), and KAPA qPCR analysis (KAPA Biosystems LIB Quant Kit, Illumina/LC480, PN: KK4854) to evaluate the quantity of sequencable transcripts.

##### Flow Cell Preparation and Sequencing

Sample concentrations are normalized to 10 nM and loaded onto Illumina Rapid flow cell at a concentration that yields 150M passing filter clusters per lane. Samples are sequenced using paired-end sequencing on an Illumina HiSeq 2500 according to Illumina protocols. The 8 bp index is read during an additional sequencing read that automatically follows the completion of read 1. Data generated during sequencing runs are simultaneously transferred to the YCGA high-performance computing cluster. A positive control (prepared bacteriophage Phi X library) provided by Illumina is spiked into every lane at a concentration of 0.3% to monitor sequencing quality in real time.

##### Data Analysis and Storage

Signal intensities are converted to individual base calls during a run using the system’s Real Time Analysis (RTA) software. Base calls are transferred from the machine’s dedicated personal computer to the Yale High Performance Computing cluster via a 1 Gigabit network mount for downstream analysis. Primary analysis – sample de-multiplexing and alignment to the human genome – is performed using Illumina’s CASAVA 1.8.2 software suite. The Cell Ranger Single-Cell Software Suite (versions 2.0.0 and 2.1.0 for the discovery and validation patients respectively) were used to perform sample demultiplexing, barcode processing and single-cell gene counting (http://10xgenomics.com/). The gene-cell matrix was generated for the following analysis.

#### Data normalization

Only genes with at least one UMI count detected in at least one cell were retained for analysis Single cells were excluded when >10% of reads mapped to mitochondrial RNA to ensure that all of the single cells originated from nucleated cells. The cells with fewer than 370 expressed genes or possible doublet cells (>100,000 reads) were also discarded. Applying these three criteria resulted in retention of 21,430 genes and 15,973 single cells for downstream analysis.

Data normalization was performed using a standard protocol in Seurat.^23^ An exploratory analysis showed cells clustered by each participant. Batch effects were then removed using an empirical Bayesian framework (ComBat function in R package sva) with the individual labeled as the random variable. The normalized data was dimensionally reduced in two dimensions using t-distributed stochastic neighbor embedding (t-SNE) with a perplexity parameter of 20 and 3000 iterations after an initial principle component analysis. The distance matrix of the single cells was computed with the output of tSNE and converted into a graph. Then, the graph was clustered with the cluster_louvain function in R package *igraph*, which implements the multi-level optimization of modularity algorithm for finding community structure.

#### Cell type identification

A gene set of the cell type markers in PBMC was manually curated from the literature. A binary cell type matrix with cell types as columns and genes as rows was generated with value 1 representing a marker in a cell type and value 0 denoting non-markers for a cell type. The generalized linear model (GLM) was constructed by deciding on one vector, representing one cell type as response, of the binary matrix and corresponding gene expressions as explanatory variables in one cell. In details, ***Y*** *= (y_gc_)_G×C_* is the binary cell type matrix of *G* marker genes of *c* cell types and ***X*** *= (x_gs_)_G×S_* is the gene expression matrix of *G* marker genes of *S* single cells. For the *g^th^* gene, *y_gc_* = 0 or 1 denotes the cell type-specific gene indicator of *c^th^* cell type and *x_gs_* denotes the gene expression of *s^th^* single cell, where *g* = 1,…, *G*, *c* = 1,…, *C*, and *s* = 1,…, *S.* The linear probability model (LPM) was conducted for our study, ***y****_c_ = β*_0*cs*_ *+ β*_1*cs*_***x****_s_* + *ε_cs_*, where ***y_c_*** = (*y*_1*c*_,…, *y_Gc_*)*^T^* and *x****_s_*** = (*x*_1*s*_,…, *x_Gs_*)*^T^* are the columns of matrix ***Y*** and ***X***, represent binary cell type vector and single cell gene expression vector, respectively. *ε_cs_ =* (*ε*_1c*s*_,…, *ε_Gcs_*)*^T^* is a random error vector, where *ε_gcs_* ∼ *N*(0, *σ*^2^). The hypothesis of interest to be tested is *H*_0*cs*_*: β*_1*cs*_ *= 0 vs H*_1*cs*_*: β*_1*cs*_ ≠ *0*. One statistical value (t value) and p value is obtained for each cell and cell type; *t_cs_* and *p_cs_* denote the test statistic value and P-value, respectively for the *c^th^* cell type of *s^th^* single cell.

We used the cutoff, p < *p*_0_ and t value > 0 to assign one cell to one cell type, where *p*_0_ is a predefined cut-off value (*p*_0_ = 0.01 in this study). Then, the proportion of each assigned cell type was calculated for every cluster. The dominant proportions are used to assign one cluster to one cell type.

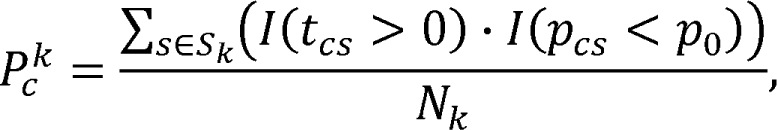

where *p_c_^k^* denotes the proportion of the *c^th^* cell type in the *k^th^* cluster. *S_k_* is the set of cell indices for clustering *k*, with *N_k_* = |*S_k_*| being the total number of cells in the *k^th^* cluster. Finally, the cell type for each cluster was confirmed manually by cell type marker gene expressions mapping on the 2D tSNE plot. The cell clusters mapping to the same cell type were merged for downstream analysis.

### Statistics

All statistical analyses were performed using R version 3.5.1 (R Foundation, https://www.r-project.org) and RStudio version 1.1.453 (https://www.rstudio.com).

#### Differential gene expression

We applied a linear mixed regression model to identify genes associated with THC infusion. To control transcriptomic variation between two participants, we used participant as a random effect. Transcriptome-wide significance was set at false discovery rate (FDR) <0.05.

#### Cell type-based gene-gene correlation analysis

The relationship among genes in each given cell type was from KEGG database. We tested whether gene links in KEGG were significant in each cell type using linear regression model. Significant gene-gene correlation was set at FDR<0.05. The analysis was performed separately in pre-THC and post-THC samples to test if THC alters gene-gene correlation.

#### Gene Ontology (GO) enrichment analysis

Genes in each co-expression network in major cell types were tested the enrichment on GO terms using The DAVID Gene Functional Classification Tool.^57^ Significant level was set at FDR <0.05.

#### Cell type-based gene set enrichment analysis

We separately analyzed gene set enrichment in each cell type using KEGG database. Differentially expressed genes in a given cell type with nominal p < 0.001 were selected. Significant pathway was set at FDR <0.05.

#### Co-expression of cannabinoid receptor genes with other genes

Four cannabinoid receptor genes, *CNR1, CNR2, CPR18*, and *GPR55* were tested differential expression across seven major cell types using linear mixed regression model. Correlation of *CNR2* with other genes in B cells was performed in pre- and post-THC samples independently. Significant correlation was set at FDR <0.05.

## Reporting summary

Further information about study design, codes, and statistics are in the reporting summery.

## Data availability

Single cell transcriptome data have been deposited in Gene Expression Omnibus and are available under project number GSE130228.

## Acknowledgement

This work was supported by the National Institute on Drug Abuse grants R01DA047820 (PI: Xu), R01DA047063 (PI: Xu), R01042691 (PI: Xu), and 1R21AA024257 (PI: D’Souza). Suhas Ganesh is partially supported by a NARSAD Young Investigator grant from the Brain and Behavior Research Foundation.

The authors thank Wenzhong Liu, Child Study Center at Yale University, Jaime Heltke, Guilin Wang, Christopher Castaldi at Yale Center of Genomic Analysis for supports of single cell RNA sequencing. The Veterans Affairs Connecticut Healthcare System (VACHS), West Haven. The nurses (Angelina Genovese, Elizabeth O’Donnell and Margaret Dion-Marovitz) of the Neurobiological studies Unit, and the research pharmacist (Rachel Galvan) of VACHS.

## Contributions

K. Xu, Y. Hu, B. Aouizerat, D.C. D’Souza, J. H. Krystal designed the study and wrote the manuscript. Y. Hu, C. Shu, X. Liang, C. Yan, analyzed data. M. Ranganathan, S. Ganesh, and D.C. D’Souza were responsible for participant recruitment and THC infusion. All authors contributed to manuscript preparation.

## Competing interests

All co-authors except Dr. Krystal declare no completing interests

The following competing interests for John H. Krystal:

**RE: John H. Krystal, MD**

**2019 Financial Disclosure**

### Consultant

Note: – The Individual Consultant Agreements listed below are less than $10,000 per year AstraZeneca Pharmaceuticals

Biogen, Idec, MA

Biomedisyn Corporation

Bionomics, Limited (Australia)

Boehringer Ingelheim International

Concert Pharmaceuticals, Inc.

Epiodyne, Inc.

Heptares Therapeutics, Limited (UK)

Janssen Research & Development

L.E.K. Consulting

Otsuka America Pharmaceutical, Inc.

Perception Neuroscience Holdings, Inc.

Spring Care, Inc.

Sunovion Pharmaceuticals, Inc.

Takeda Industries

Taisho Pharmaceutical Co., Ltd

### Scientific Advisory Board

Bioasis Technologies, Inc.

Biohaven Pharmaceuticals

BioXcel Therapeutics, Inc. (Clinical Advisory Board)

Cadent Therapeutics (Clinical Advisory Board)

PsychoGenics, Inc.

Stanley Center for Psychiatric research at the Broad Institute of MIT and Harvard

Lohocla Research Corporation

### Stock

ArRETT Neuroscience, Inc.

Biohaven Pharmaceuticals

Sage Pharmaceuticals

Spring Care, Inc.

### Stock Options

Biohaven Pharmaceuticals Medical Sciences

BlackThorn Therapeutics, Inc.

Storm Biosciences, Inc.

### Income Greater than $10,000

### Editorial Board

Editor – Biological Psychiatry

### Patents and Inventions

1. Seibyl JP, Krystal JH, Charney DS. Dopamine and noradrenergic reuptake inhibitors in treatment of schizophrenia. US Patent #:5,447,948.September 5, 1995
2. Vladimir, Coric, Krystal, John H, Sanacora, Gerard – Glutamate Modulating Agents in the Treatment of Mental Disorders US Patent No. 8,778,979 B2 Patent Issue Date: July 15, 2014. US Patent Application No. 15/695,164: Filing Date: 09/05/2017
3. Charney D, Krystal JH, Manji H, Matthew S, Zarate C., – Intranasal Administration of Ketamine to Treat Depression United States Application No. 14/197,767 filed on March 5, 2014; United States application or Patent Cooperation Treaty (PCT) International application No. 14/306,382 filed on June 17, 2014
4. Zarate, C, Charney, DS, Manji, HK, Mathew, Sanjay J, Krystal, JH, Department of Veterans Affairs “Methods for Treating Suicidal Ideation”, Patent Application No. 14/197.767 filed on March 5, 2014 by Yale University Office of Cooperative Research
5. Arias A, Petrakis I, Krystal JH. – Composition and methods to treat addiction. Provisional Use Patent Application no.61/973/961. April 2, 2014. Filed by Yale University Office of Cooperative Research.
6. Chekroud, A., Gueorguieva, R., & Krystal, JH. “Treatment Selection for Major Depressive Disorder” [filing date 3rd June 2016, USPTO docket number Y0087.70116US00]. Provisional patent submission by Yale University
7. Gihyun, Yoon, Petrakis I, Krystal JH – Compounds, Compositions and Methods for Treating or Preventing Depression and Other Diseases. U. S. Provisional Patent Application No. 62/444,552, filed on January10, 2017 by Yale University Office of Cooperative Research OCR 7088 US01
8. Abdallah, C, Krystal, JH, Duman, R, Sanacora, G. Combination Therapy for Treating or Preventing Depression or Other Mood Diseases. U.S. Provisional Patent Application No. 047162-7177P1 (00754) filed on August 20, 2018 by Yale University Office of Cooperative Research OCR 7451 US01

### NON-Federal Research Support

AstraZeneca Pharmaceuticals provides the drug, Saracatinib, for research related to NIAAA grant “Center for Translational Neuroscience of Alcoholism [CTNA-4]

## Supplementary Figures

**Figure S1. tSNE plot showing cell clusters after removed batch effects**

a. tSNE plot for four samples of pre- and post-THC infusion.
b. tSNE plot for pre-THC infusion of subject 1
c. tSNE plot for post-THC infusion of subject 1
d. tSNE plot for pre-THC infusion of subject 2
e. tSNE for post-THC infusion of subject 2.

**Figure S2. Sample marker gene expression in each cell type of 15,973 peripheral blood mononuclear cells**

**Figure S3. Study design and procedure of THC infusion in human subjects.** SA: Substance Abuse; PANSS: Positive and Negative Syndrome Scale; CADSS: Clinician Administered Dissociative States Scale; VAS: Visual Analog Scale; IV: Intravenous Injection; EEG: electroencephalogram.

## Supplementary Tables

**Table S1.** Cell markers for cell type identification

**Table S2.** Mapping cell types using generalized linear modelling

**Table S3.** Proportion of cell numbers for each sample in a given cell type

**Table S4.** A summary of the number of differentially expressed genes between post- and pre-THC samples in each cell type

**Table S5.** Differentially expressed genes affected by THC in CD4+ T-cells

**Table S6.** Differentially expressed genes affected by THC in IL7RCD4+ T-cells

**Table S7.** Differentially expressed genes affected by THC in CD8+ T-cells

**Table S8.** Differentially expressed genes affected by THC in B cells

**Table S9.** Differentially expressed genes affected by THC in natural killer cells

**Table S10.** Differentially expressed genes affected by THC in CD14+ monocytes

**Table S11.** Differentially expressed genes affected by THC in FCGR3A monocytes

**Table S12.** log2 fold change of top 20 genes between post- and pre-THC infusion in each CD4+ T-cell clusters

**Table S13**. log2 fold change of top 20 genes between post- and pre-THC infusion in each CD8+ T-cell clusters

**Table S14.** log2 fold change of top 20 genes between post- and pre-THC infusion in each B cell clusters

**Table S15.** log2 fold change of top 20 genes between post- and pre-THC infusion in each CD14+ monocytes clusters

**Table S16.** Gene Ontology (GO) term enrichment for the co-expression network in CD4+ T-cells

**Table S17.** Gene Ontology (GO) term enrichment for the co-expression network in CD8+ T-cells

**Table S18.** Gene Ontology (GO) term enrichment for the co-expression network in B cells

**Table S19.** Gene Ontology (GO) term enrichment for the co-expression network in natural killer cells

**Table S20.** Gene Ontology (GO) term enrichment for the co-expression network in CD14+ monocytes

**Table S21.** Gene Ontology (GO) term enrichment for the co-expression network in FCGR3A monocytes

**Table S22.** A summary of the number of significant pathway in each cell type

**Table S23.** Significant pathways from KEGG in CD4+ T-cells

**Table S24.** Significant pathways from KEGG in IL7RCD4+ T-cells

**Table S25.** Significant pathways from KEGG in CD8+ T-cells

**Table S26.** Significant pathways from KEGG in B cells

**Table S27.** Significant pathways from KEGG in natural killer cells

**Table S28.** Significant pathways from KEGG in CD14+ monocytes

**Table S29.** Significant pathways from KEGG in FCGR3A monocytes

**Table S30.** Pre-THC gene co-expression between *CNR2* and other genes

**Table S31.** Post-THC gene co-expression between *CNR2* and other genes

