## Supplementary figures and images for "Single-cell Transcriptome Mapping Identifies Common and Cell-type Specific Genes Affected by Acute Delta9-tetrahydrocannabinol in Humans"

### Figure S1a

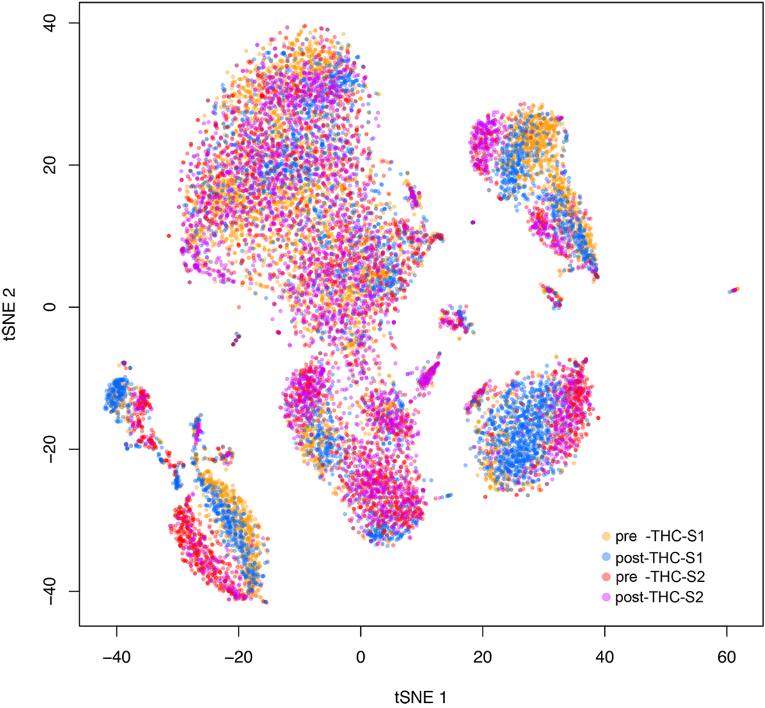

### Figure S1b

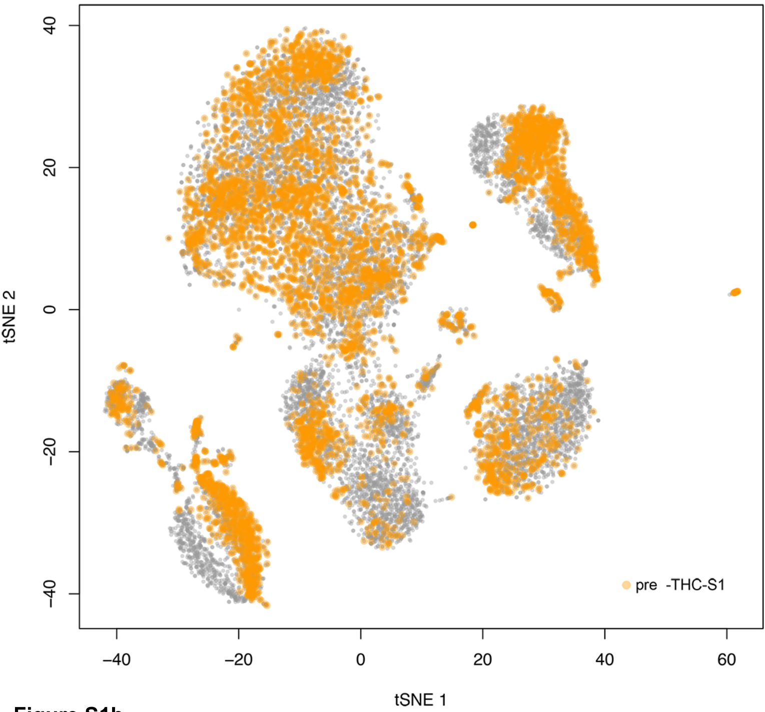


**Figure S1b**

### Figure S1c

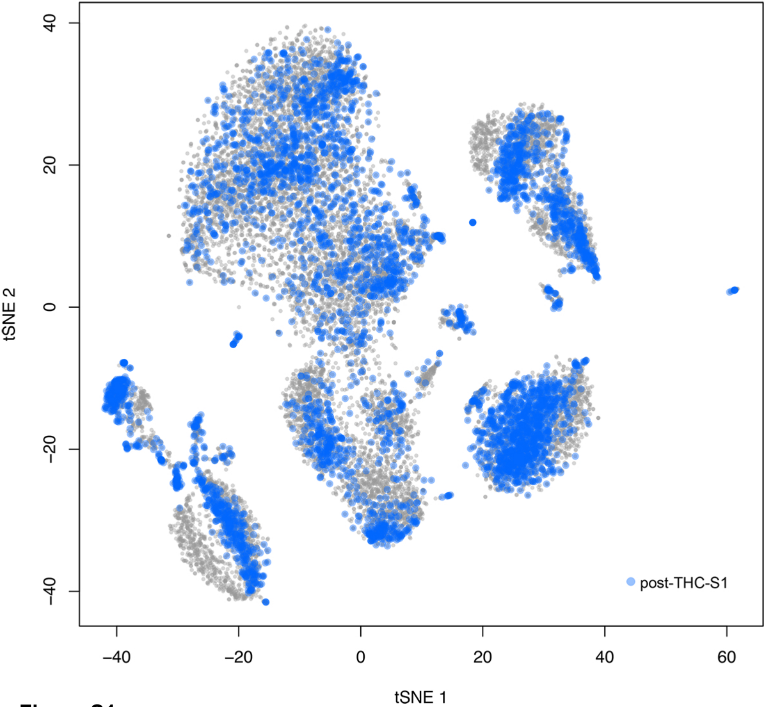


**Figure S1c**

### Figure S1d

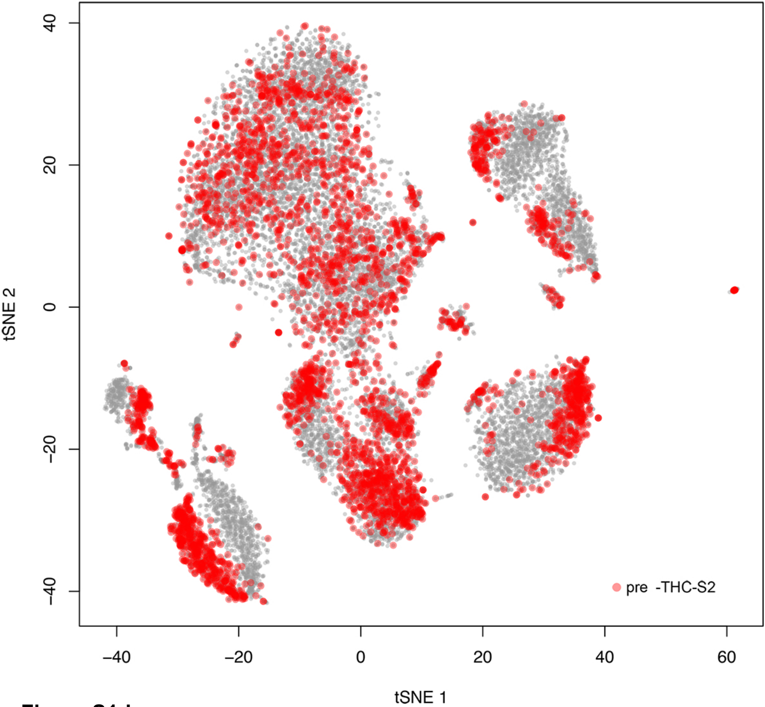


**Figure S1d**

### Figure S1e

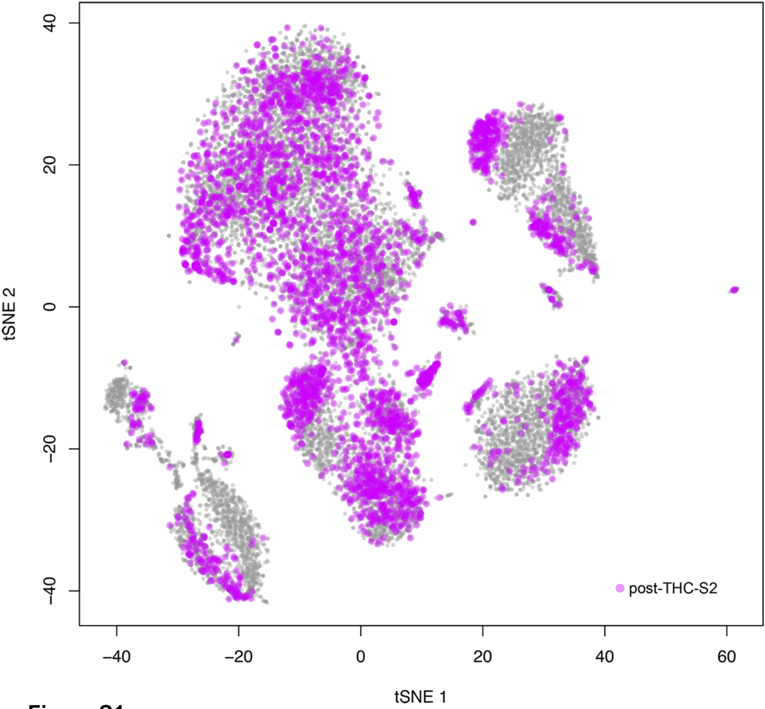


**Figure S1e**

### Figure S2

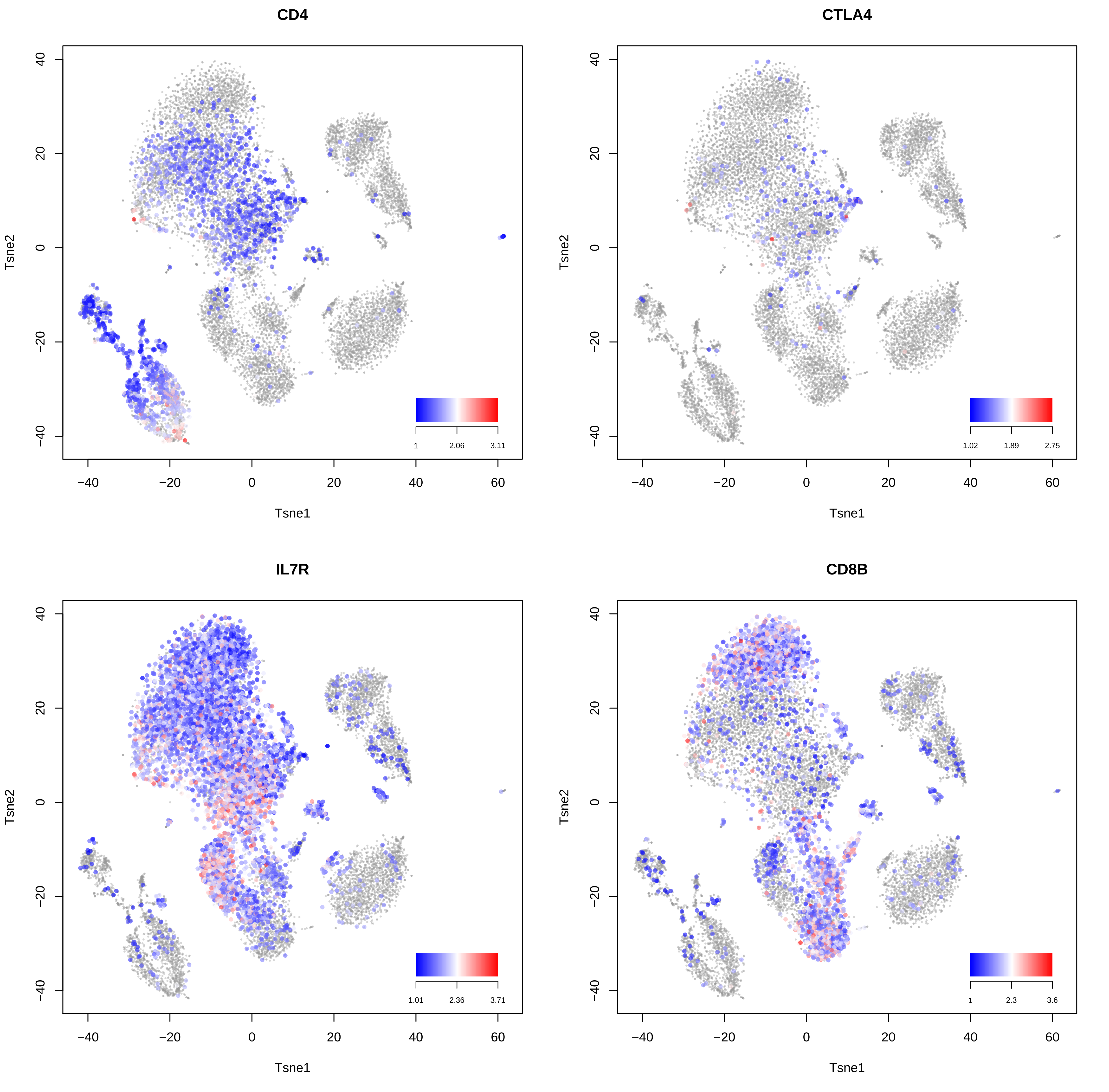

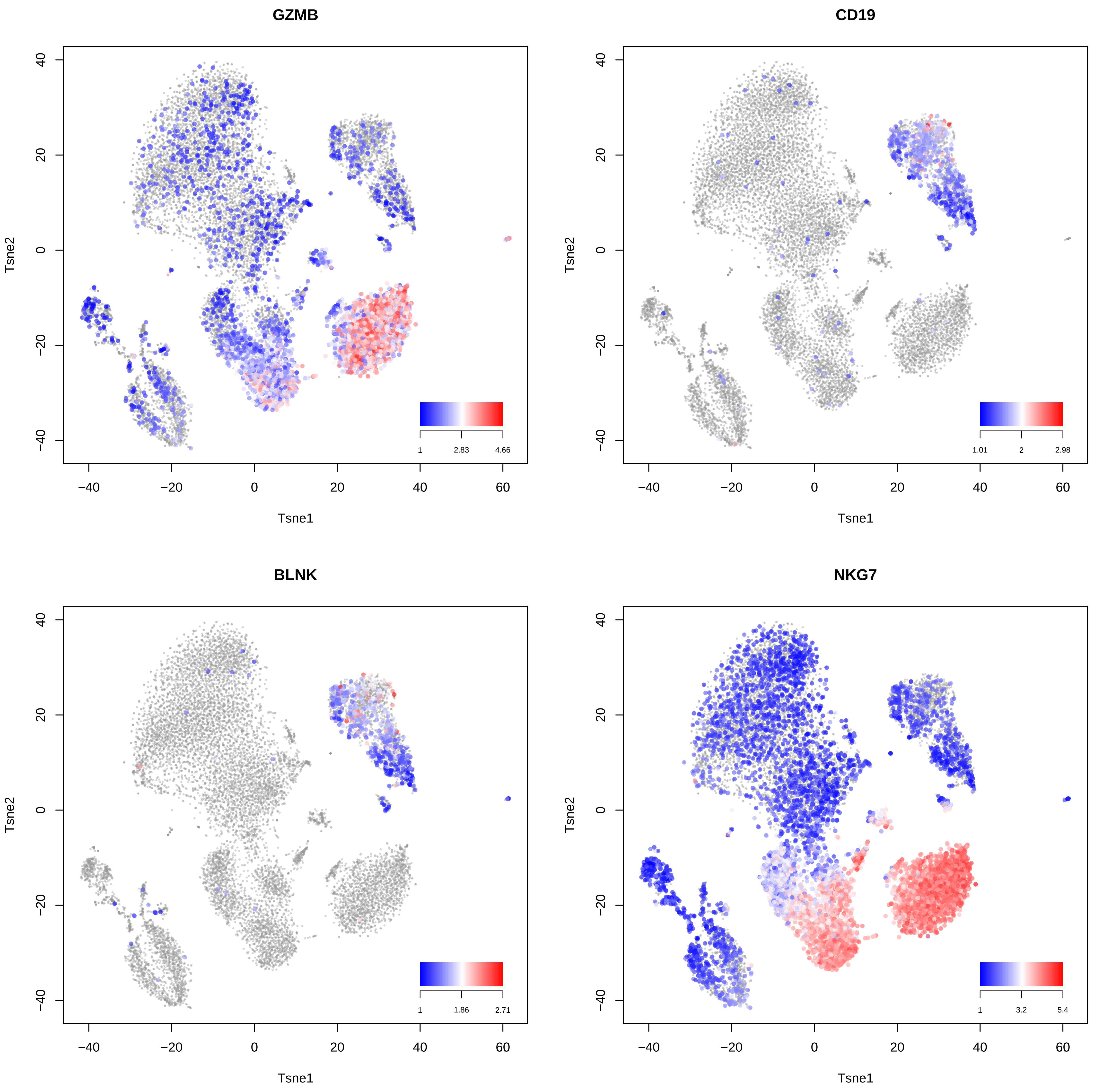

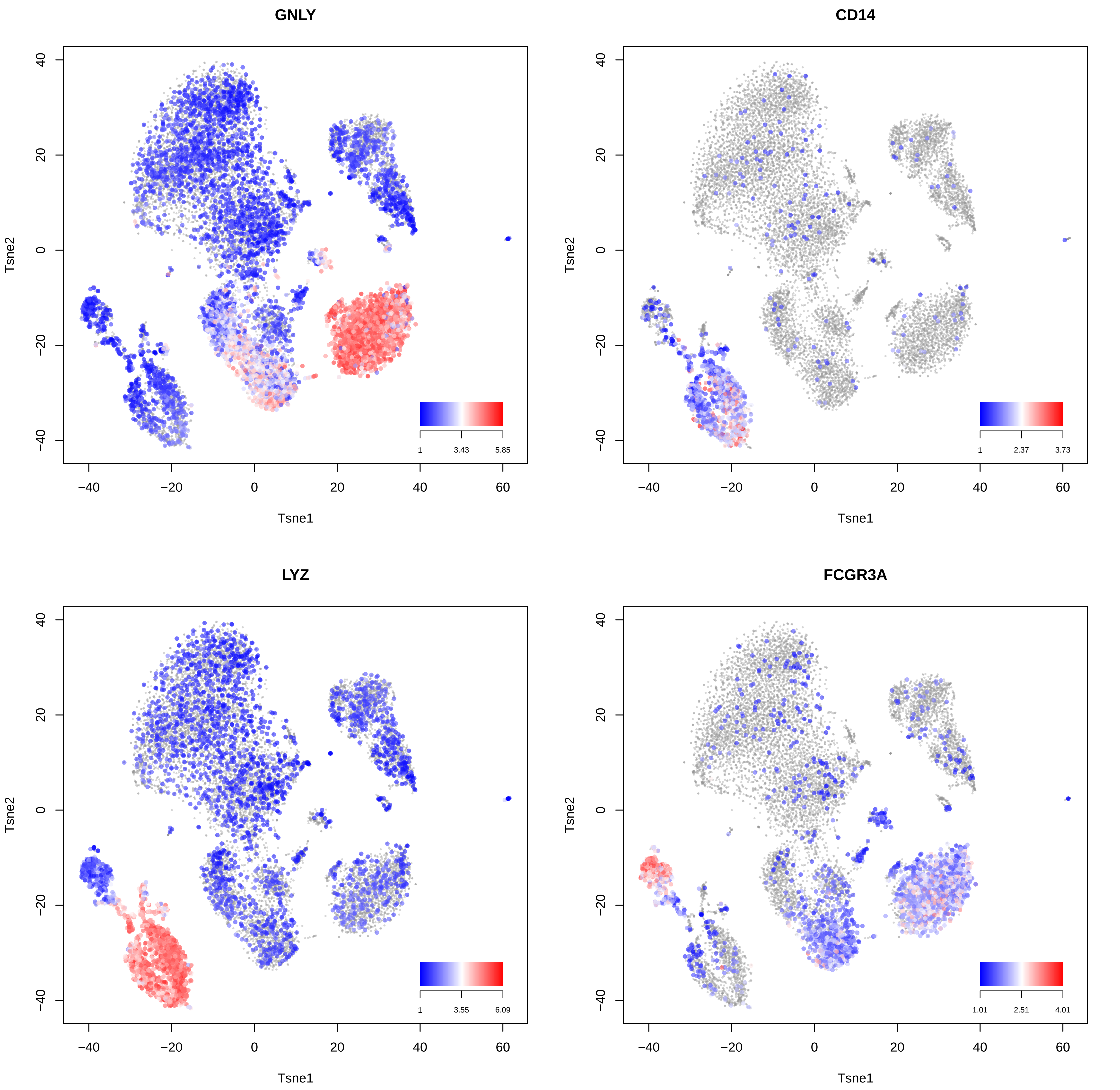

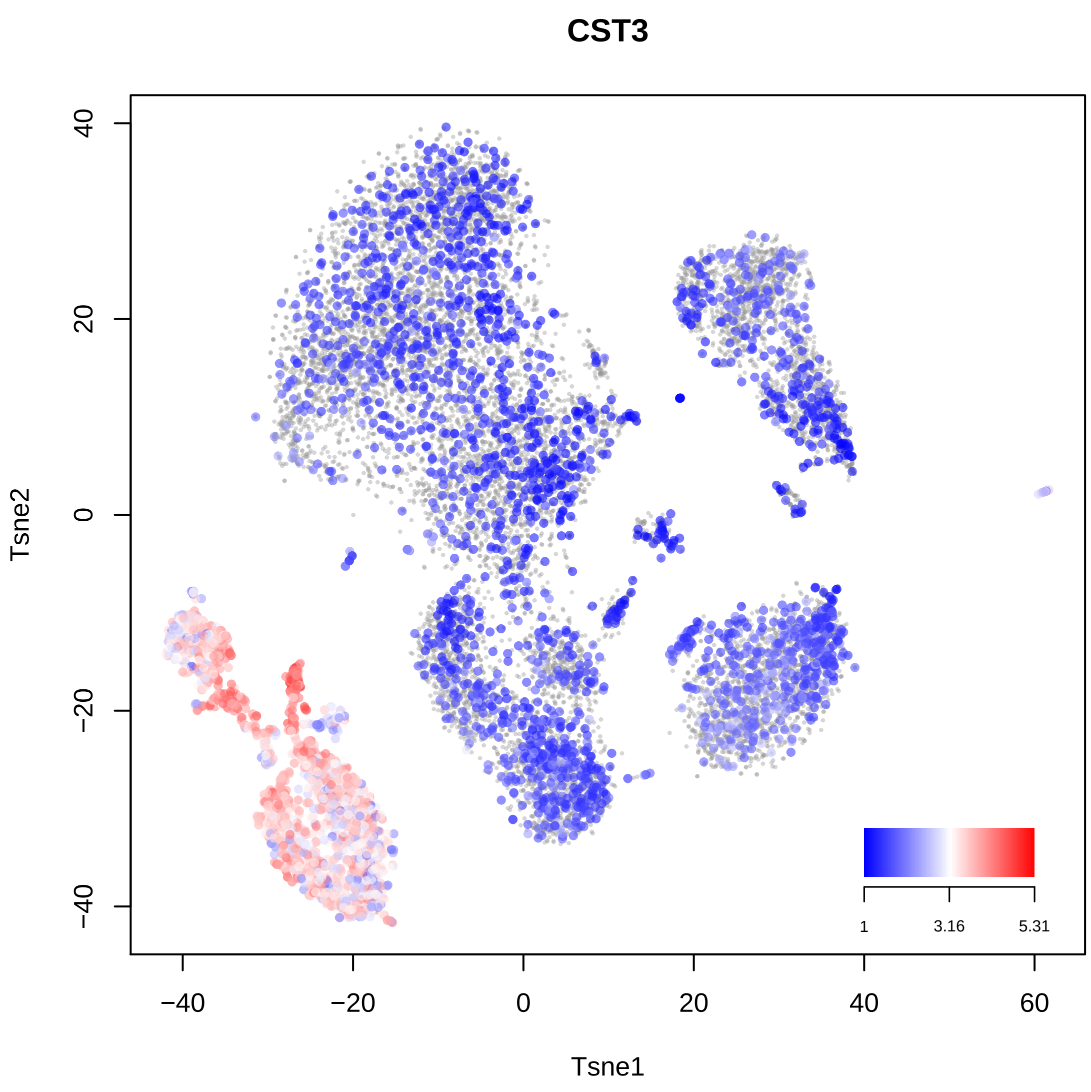

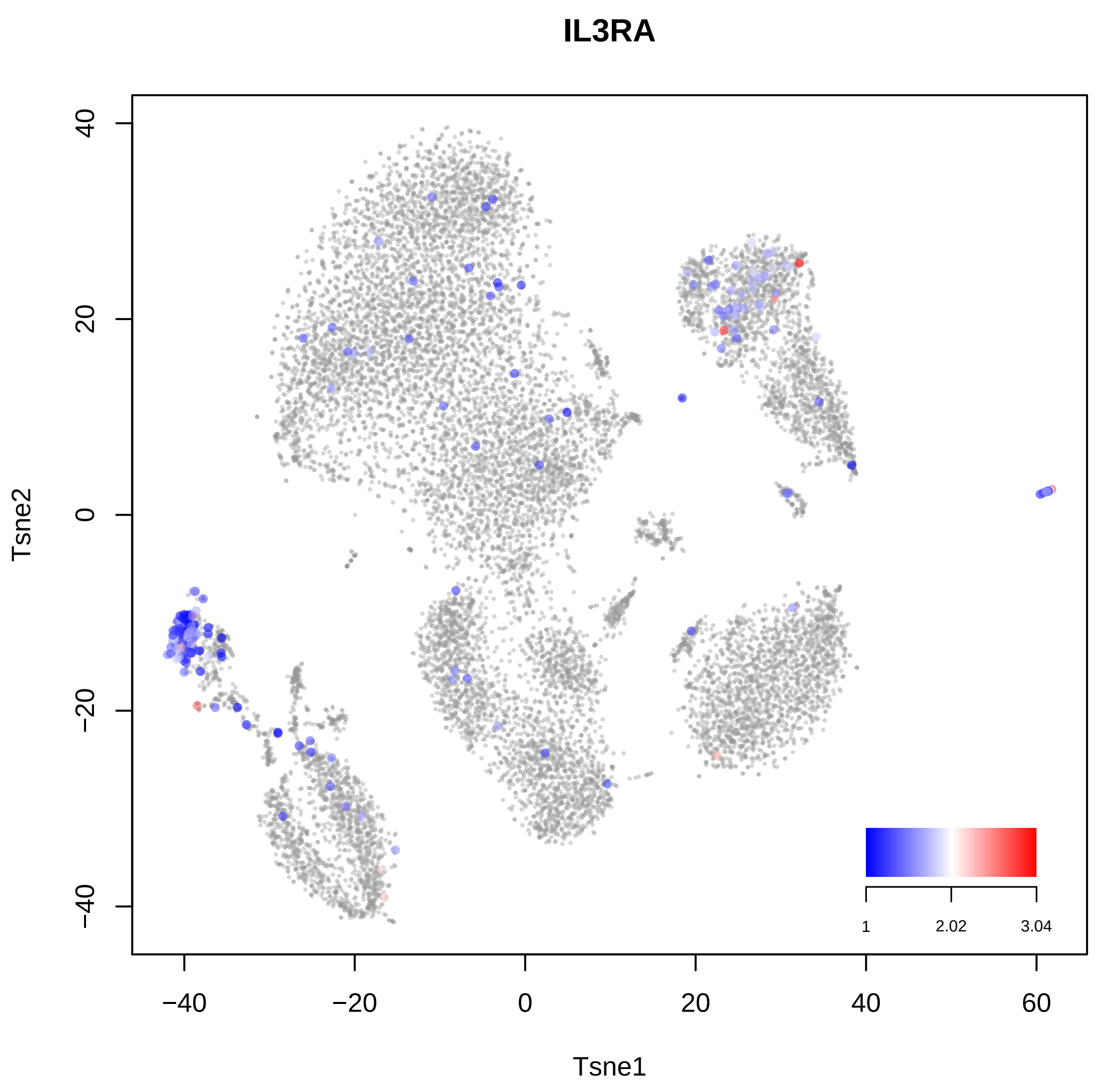

### Figure S3

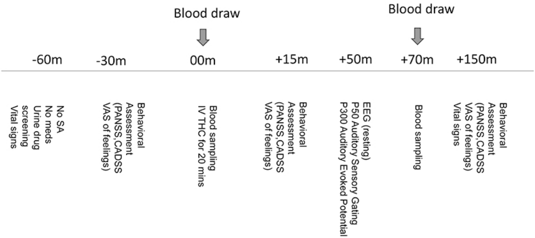


**Figure S3**. Study design and procedure
